# Unblended Disjoint Tree Merging using GTM improves species tree estimation

**DOI:** 10.1101/835959

**Authors:** Vladimir Smirnov, Tandy Warnow

**Affiliations:** University of Illinois at Urbana-Champaign

## Abstract

Phylogeny estimation is an important part of much biological research, but large-scale tree estimation is infeasible using standard methods due to computational issues. Recently, an approach to large-scale phylogeny has been proposed that divides a set of species into disjoint subsets, computes trees on the subsets, and then merges the trees together using a computed matrix of pairwise distances between the species. The novel component of these approaches is the last step: Disjoint Tree Merger (DTM) methods. We present GTM (Guide Tree Merger), a polynomial time DTM method that adds edges to connect the subset trees, so as to provably minimize the topological distance to a computed guide tree. Thus, GTM performs *unblended* mergers, unlike the previous DTM methods. Yet, despite the potential limitation, our study shows that GTM has excellent accuracy, generally matching or improving on two previous DTMs, and is much faster than both. Thus, the GTM approach to the DTM problem is a useful new tool for large-scale phylogenomic analysis, and shows the surprising potential for unblended DTM methods. The software for GTM is available at https://github.com/vlasmirnov/GTM.

## 1 Background

The estimation of evolutionary trees (i.e., phylogenies, whether of genes or of species) is a fundamental step in much biological research, including understanding how species adapt to their environments, how gene function evolves, and how humans migrated across the globe. Yet, phylogeny estimation is computationally intensive, as nearly all the best approaches are based on NP-hard optimization problems.

Divide-and-conquer approaches to tree estimation have the potential to provide improved scalability to heuristics for NP-hard optimization problems, but they depend on supertree methods, which have not been established to be highly scalable [1]. A new divide-and-conquer approach has been recently proposed to tackle this limitation: the set of species is divided into smaller, disjoint subsets, trees are computed on each subset (using the best available methods), and then the trees are combined together into a tree on the complete taxon set. In this approach, the subset trees are considered hard constraints and so must be induced in the output tree [2, 3, 4, 5, 6]; this property of the output tree is expressed by saying it is a “compatibility supertree” of the input constraint trees. Finally, merging disjoint trees requires some auxiliary information (such as a matrix of pairwise distances between the species), and the problem of merging disjoint trees using auxiliary information is called the “Disjoint Tree Merger” (DTM) problem.

So far, three DTM methods have been developed: NJMerge [2, 3], TreeMerge [4], and constrained-INC [6, 5], each of which uses a computed matrix of pairwise distances to merge the disjoint trees and does so in polynomial time. Constrained-INC was tested only for gene tree estimation, where it had disappointing results. TreeMerge and NJMerge were tested in the context of species tree estimation on multi-locus datasets, where they provided good advantages: they matched the accuracy of leading species tree estimation methods (ASTRAL [7, 8, 9] and RAxML [10], which performs a concatenation maximum likelihood analysis), while providing improvements in speed. NJMerge can fail to return a tree under some conditions (though never when merging two trees), due to its algorithmic design [2, 3]; TreeMerge is a technique that uses NJMerge on pairs of trees (where it is guaranteed to never fail), and then combines overlapping trees using an innovative algorithm that exploits the ability to estimate branch lengths. Thus, TreeMerge was developed specifically to replace NJMerge because of the capacity of NJMerge to fail and in order to improve the speed. Finally, all three DTM methods have been proven to enable statistically consistent tree estimation when used within appropriate divide-and-conquer strategies.

We propose a new DTM approach, where the input is a set of disjoint trees and also a tree on the full set of species (called a “guide tree”), and the objective is to produce a tree that agrees with all constraint trees and minimizes the total topological distance to the guide tree. We prove this optimization problem is NP-hard, but show that if we do not allow blending (so that the output tree is formed by just adding edges between the constraint trees), then the problem is solvable in polynomial time.

We present the Guide Tree Merger (GTM) algorithm to solve this unblended DTM-GT problem, and prove that it uses polynomial time. We evaluate GTM for species tree estimation from multi-locus datasets, where gene trees can differ from the species tree due to incomplete lineage sorting [11]. We prove that the pipeline maintains statistical consistency, and demonstrate (on a collection of multi-locus datasets with 1000 species) that GTM has very good performance. Specifically, we show that GTM matches or improves on the accuracy of NJMerge and TreeMerge on these datasets, while completing in a fraction of their runtimes.

## 2 The DTM-GT optimization problem

All phylogenetic trees are assumed to be unrooted and leaf-labelled by taxa in a set *S*. We continue with some basic terminology.

Given a tree *T* on leafset *S* and set *S*′ ⊆ *S*, the homeomorphic subtree of *T* defined by *S*′ (in which degree two nodes are suppressed) is denoted by *T* |_*S*′_.

### Definition 1.

*Let* 𝒜 = {*T*_1_, *T*_2_, …, *T*_*k*_} *be a set of trees on subsets of S. T is said to be a* ***compatibility supertree*** *for* 𝒜 *if (1) L*(*T*) = ∪_*i*_*L*(*T*_*i*_) *(where L*(*t*) *denotes the leaf set of t) and (2) T* |_*L*(*t*)_ = *t for all t* ∈ 𝒜.

Each edge *e* in a phylogenetic tree *t* defines a bipartition *π*_*e*_ on the leafset of *t*, and hence each tree *t* is defined by the set of its bipartitions, which we denote by *C*(*t*) = {*π*_*e*_|*e* ∈ *E*(*T*)}. The “FN” (false negative) distance of tree *A* to tree *B* is |*C*(*B*)\*C*(*A*)|, or the number of bipartitions in *C*(*B*) that are missing from *C*(*A*), and is denoted by *FN* (*A, B*).

We continue with a discussion of DTM methods. Each DTM method takes as input a set of 𝒯 leaf-disjoint trees and some auxiliary information and returns a compatibility supertree for 𝒯. Note that since the trees in 𝒯 are on disjoint leaf sets, a compatibility supertree is guaranteed to exist (e.g., the tree formed by including a center node *v* and making *v* adjacent to some internal vertex in each of the trees in 𝒯 is a compatibility supertree). The key to making DTM methods have good accuracy is the use of auxiliary information to merge the trees together well.

The current DTM methods perform this merger by computing a matrix of pairwise distances between species under a statistical model of evolution, and then carefully use the resultant matrix while merging the subset trees. For example, NJMerge accomplishes this by modifying the popular Neighbor Joining method [12], which agglomeratively builds the tree, so that it never violates any subset tree in 𝒯. Furthermore, the current DTM methods allow blending, by which we mean that the subset trees in 𝒯 may not be separated from each other within the compatibility supertree.

To understand why blending can be necessary, suppose *D* is an additive matrix (i.e., there is an edge-weighted tree *T* so that *D*[*x, y*] is the distance between *x* and *y* in *T*) that defines a caterpillar tree on 8 leaves, i.e., *T* = (1, (2, (3, (4, (5, (6, (7, 8))))))). Now suppose we have two trees, *t*_1_ and *t*_2_, with *t*_1_ = (1, (5, (6, 7))) and *t*_2_ = (2, (3, (4, 8))). Now consider when the input to the DTM method is the pair 𝒯 = {*t*_1_, *t*_2_} and *D*. The desired output should be *T*, which is a compatibility supertree for 𝒯 that realizes the input matrix *D*. Yet, *T* can only be formed if *t*_1_ and *t*_2_ are *blended* together; thus, unblended merges (formed by adding an edge connecting the input trees) are not as powerful as blended mergers.

In this paper, we introduce GTM (Guide Tree Merger), a new DTM method that does not allow blending. Despite this limitation, we will show GTM matches or improves on the previous DTM methods NJMerge and TreeMerge under most tested conditions, and is much faster than both. Before we show our method, we introduce an optimization problem for DTM construction.

### Definition 2. DTM-GT-FN

- *Input: A set 𝒯* = {*T*_1_, *T*_2_, …, *T*_*k*_} *of leaf-disjoint unrooted binary trees and a guide tree T* * *on the full set of species*
- *Output: A compatibility supertree T for 𝒯 that minimizes* |*C*(*T* *) \ *C*(*T*)| *(i.e., T minimizes the FN distance to T* *).

Here, GT refers to “guide tree” and “FN” refers to the optimization criterion, which is that we wish to minimize the FN (false negative) distance between the guide tree and the compatibility supertree we construct. We allow the trees in the input set 𝒯 to be non-binary, and for this reason we use the FN distance rather than the RF distance. However, when the guide tree and all the trees in 𝒯 are binary, then the optimal compatibility supertree is also binary, and the RF distance and FN distance to the guide tree are identical.

### Theorem 1. DTM-GT-FN is NP-hard

See Appendix B for the proof. Now consider the following variant of DTM-GT-FN where we do not permit blending:

### Definition 3. Unblended DTM-GT-FN

- *Input: A set 𝒯* = {*T*_1_, *T*_2_, …, *T*_*k*_} *of leaf-disjoint unrooted binary trees and a guide tree T* * *on the full set of species*
- *Output: A compatibility supertree T for 𝒯 formed by connecting the trees in 𝒯 by edges and that minimizes* |*C*(*T* *) \ *C*(*T*)| *(i.e., the FN distance to T* \**) among all such compatibility supertrees.*

Interestingly, Unblended-DTM-GT-FN can be solved in polynomial time, as we show in the next section.

## 3 The GTM method

We begin with some terminology. First, we say that a bipartition *π* = *A*|*B* “violates a constraint tree” *t* if *t* does not contain any edge defining the bipartition *π*|_*L*(*t*)_ = *A* ∩ *L*(*t*)|*B* ∩ *L*(*t*), where *π*|_*L*(*t*)_ denotes *π* restricted to *L*(*t*). Next, we refer to the leaf sets of each constraint tree as “constraint sets”, and we partition the edges of the guide tree *T* * into three sets, based on how many constraint sets (0, 1, or at least 2) have leaves on both sides of the edge.

- The edges that have no constraint sets on both sides are called “bridge edges”, as they separate constraint sets.
- The edges that have exactly one constraint set on the two sides are said to “lie within that constraint set.”
- The edges that have two or more constraint sets on the two sides are said to “violate convexity” (where convexity would assert that the nodes of the guide tree can be labelled by the constraint sets, so that no node has more than one label and every two leaves with the same label are connected by a path with all nodes given the same label).

GTM solves the Unblended DTM-GT-FN problem recursively, in polynomial time, as described below. Specifically, GTM uses the guide tree to determine which pairs of constraint trees should be connected directly by edges, and then uses bipartitions in the guide tree to determine how to connect the constraint trees together; this depends on finding appropriate edges in each constraint tree to subdivide (by adding internal nodes) and then which pairs of constraint trees should be connected by new edges (introduced between the newly created internal nodes in the subdivided edges). The details of how these edges are identified and how the specific pairs of constraint trees are merged are non-trivial, and are described below:

The GTM algorithm has the following structure:

- Collapse all edges in *T* * that violate any *T*_*i*_.
- Collapse all edges in *T* * that violate convexity.
- Return *Rejoin*(*T* *).

Note that *Rejoin*(·) is run only after all edges that violate convexity or a constraint tree have been collapsed. Hence, every remaining edge is either a bridge edge (i.e., separates constraint sets) or is on a path between two leaves for exactly one constraint subset (i.e., “lies within” their constraint sets). Furthermore, every edge *e* that lies within constraint set *L*(*T*_*i*_) satisfies *b*(*e*): = *π*(*e*)|_*L*(*Ti*_) ∈*C*(*T*_*i*_) (as otherwise it would have violated the constraint tree and would have been collapsed). Given this context, we now describe the recursive function *Rejoin*(*T*) (also see Figure 2):

**Figure 1:**
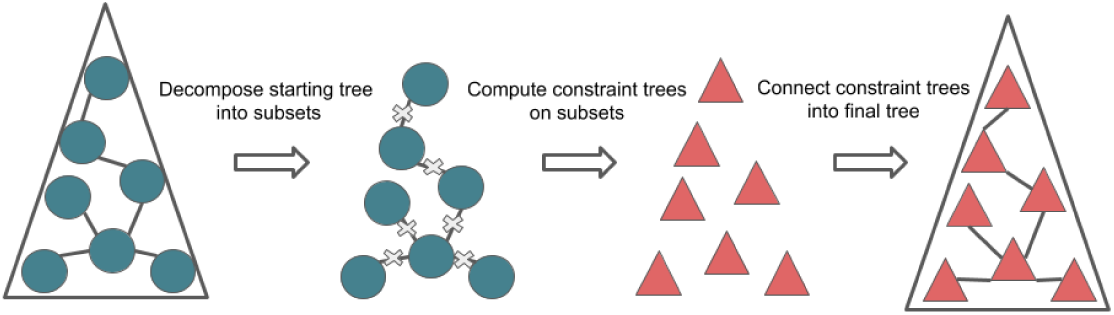
A basic DTM pipeline with GTM. A starting tree is computed and then decomposed by deleting some set of edges. Constraint trees are computed for these subsets and attached together with new edges using GTM.

**Figure 2:**
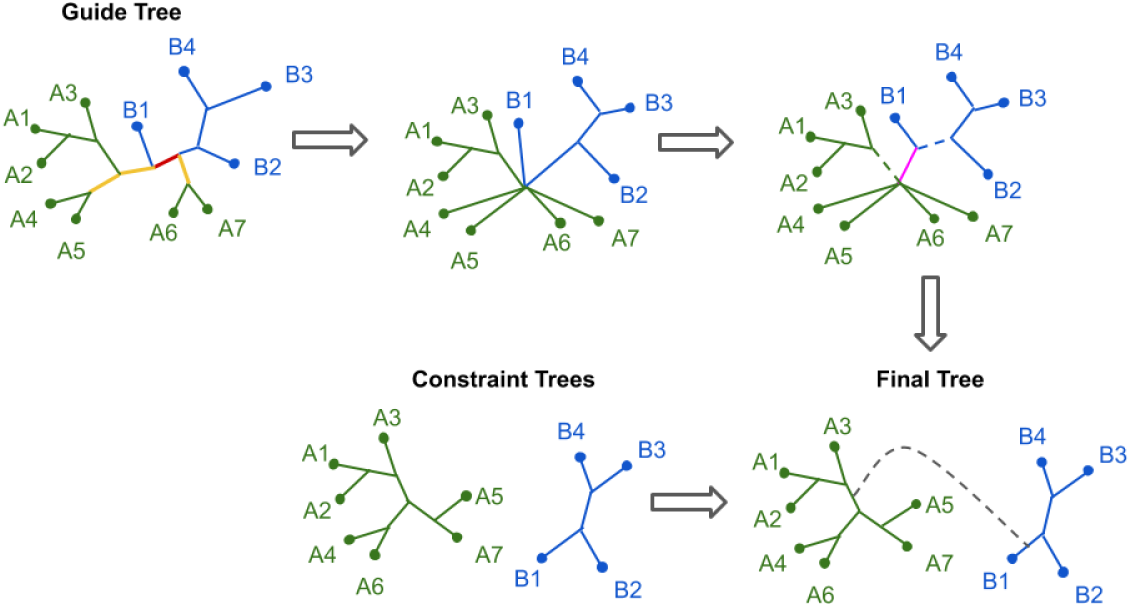
GTM example with two constraint trees (or one recursive step). Green and blue edges lie inside different constraint sets. (1) We first collapse all edges that violate convexity (red) or violate a constraint tree (yellow). (2) At the polytomy where green and blue edges meet, we separate the green and blue edges with bridge *e* (pink). (3) We randomly select one edge (dashed) at each endpoint of the bridge. (4) We locate the edges in the constraint trees that induce the same bipartition, when restricted to their respective constraint set (in this case -*A*_1_*A*_2_*A*_3_ |*A*_4_*A*_5_*A*_6_*A*_7_ and *B*_1_ |*B*_2_*B*_3_*B*_4_). (5) We join the constraint trees by subdividing the two identified edges (i.e., by adding a new internal node into each edge) and add the edge between the newly created internal nodes.

*Rejoin*(*T*):

- If *L*(*T*) = *L*(*T*_*i*_) for some *i*, Return *T*_*i*_.
- Pick an edge *e* in *T* as follows. Let *v* ∈ *T* be any “border node”, i.e., a node that is either (1) an endpoint of a bridge edge, or (2) a common endpoint of two edges lying in different constraint sets *L*(*T*_*i*_) and *L*(*T*_*j*_). Given *v*, we define edge *e* as follows: in case (1), *e* is the bridge edge that *v* is incident with, and in case (2), we refine at node *v* by adding a vertex *v*′ and new edge (*v, v*′) to separate all the edges incident with *v* that lie in *L*(*T*_*i*_) from all the other edges, and set *e* = (*v, v*′).
- Delete *e* from *T* to form two trees, *S*_1_ and *S*_2_. /*Comment: The species in each constraint subset are in exactly one subtree, *S*_1_ or *S*_2_. */
- Let edges *e*_1_ ∈ *E*(*S*_1_) and *e*_2_ ∈ *E*(*S*_2_) be any two edges that are incident with *e*.
- Let 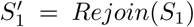 and 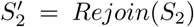. /*Comment: Note that 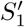 and 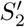 are both unblended compatibility supertrees of some of the constraint trees, and each constraint tree appears within one of these two trees. */
- Find “attachment edges” 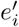 in trees 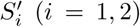, such that *e*_*i*_ and 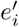 define the same bipartitions in their respective trees.
- Join 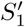 and 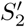 by “adding an edge between 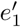 and 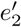” (i.e., by subdividing 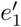 and 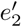, thus creating new nodes *v*_1_ and *v*_2_ and adding edge (*v*_1_, *v*_2_)) and return the resulting tree.

In other words, for each pair of constraint trees to be joined, we add an edge to connect them by choosing one edge from each tree to bridge together. This “attachment edge” is chosen by finding an edge with the same bipartition, relative to the subtree, as any of the edges at the attachment point of that subtree in *T* *.

### Theorem 2.

*Given a guide tree T* * *and set* T = {*T*_1_, *T*_2_, …, *T*_*k*_} *of constraint trees over their respective leaf sets L*(*T*_*i*_) *of* 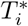, *GTM optimally solves Unblended DTM-GT-FN on* (T, *T* *), *and has a worst case running time complexity of O*(*N*^2^), *where N* = |*L*(*T* *)| *is the total number of species.*

We sketch the proof here, and direct the reader to Appendix B for full details. We note that another way of describing GTM is that it collapses all edges in the guide tree that violate convexity or violate a constraint tree, it adds edges to separate the constraint sets, and then it refines the resultant tree to induce each of the constraint trees. As a result, it is easy to see that it maximizes the number of shared bipartitions, and so equivalently it minimizes the number of missing branches in *T* *. Hence, it optimally solves Unblended DTM-GT-FN. For the running time, we add that to achieve the *O*(*N*^2^) time complexity, the algorithm uses hashing.

## 4 Divide-and-conquer pipelines for DTMs

Here we describe a generalization of the divide-and-conquer pipelines that were previously used to evaluate DTMs in [2, 3, 4, 6] in the context of multi-locus species tree estimation and gene tree estimation.

- Construct a starting tree *T*_0_ using a fast but statistically consistent method, *X*
- Divide the species set into disjoint sets by decomposing the tree *T*_0_ (by deleting edges) until each subset has at most *k* species
- Construct trees on each subset using the preferred method, *Y*
- Combine the subset trees together using a DTM method, *Z*

We will refer to a given pipeline for multi-locus species tree estimation as the triple “*X*-*Y* -*Z*”; for example “NJst-RAxML-TreeMerge” indicates that NJst [13] is used to compute the starting tree, RAxML is used to construct subset trees, and TreeMerge is used to combine the subset trees into one tree. In our experiments, we repeatedly delete centroid edges from the starting tree until all the subsets have at most 120 species (see Appendix A).

A desirable property of a phylogeny estimation method or pipeline is that it is statistically consistent, which means that it provably converges to the true tree as the amount of data increases. Pipelines using NJMerge and TreeMerge were proven statistically consistent under the MSC+GTR model [3, 4] where gene trees evolve within the species tree under the multi-species coalescent (MSC) model [14] and then sequences evolve down each gene under the Generalized Time Reversible (GTR) model [15]) and a similar pipeline using constrained-INC was proven to be statistically consistent under the GTR model [5]. The following theorem shows that appropriate divide-and-conquer pipelines using GTM are also statistically consistent; see Appendix B for the proof.

### Theorem 3.

*Let* Φ *be a model of evolution. If the method X used to construct the starting tree and the method Y used to construct the subset trees are both statistically consistent under* Φ, *then the DTM pipeline X-Y -GTM is also statistically consistent under* Φ.

Hence, NJst-ASTRAL-GTM is statistically consistent under the MSC+GTR model, but RAxML, FastTree2 [16], and any divide-and-conquer strategies based on these methods will not be (because maximum likelihood is not statistically consistent under the MSC+GTR model [17, 18]). Note that statistical consistency does not depend on using the centroid edge decomposition, and instead is guaranteed for any decomposition strategy that operates by removing a set of edges from the starting tree and computing subset trees on the leaf sets of the resulting components.

The terms “starting tree” and “guide tree” can have somewhat different meanings in divide-and-conquer pipelines. The starting tree is used to produce subsets, while the guide tree is part of the input to the GTM algorithm. Within our “*X*-*Y* -GTM” pipelines, the starting tree (*X*) and the guide tree are always the same, permitting the terms to be used interchangeably in this context. However, other pipelines can be considered where the decomposition into subsets is based on a starting tree and then the subset trees are merged using a guide tree that is different from the starting tree.

## 5 Experimental Study

We give an overview of the experimental study here; see Appendix A for additional details and software commands used in this study.

### 5.1 Overview

We compare GTM to NJMerge and TreeMerge with respect to species tree error using previously published multi-locus datasets with 1000 species and varying numbers of genes where gene trees can differ from the true species tree due to incomplete lineage sorting. We used multi-locus datasets and estimated gene trees (computed using FastTree2) from [3].

We vary the starting tree and method to compute constraint trees, so as to explore the impact of these choices on accuracy of the final tree. We compare each estimated species tree to the true species tree, and report the normalized Robinson-Foulds (RF) error rates [19], where the RF error of tree *T* is the total number of bipartitions that appear in *T* or the true tree but not both (RF distance), divided by 2*N*−6, where *N* is the number of leaves (note that 2*N*−6 is the total number of non-trivial bipartitions if both trees are binary). Thus, the RF error ranges from 0 (perfect match) to 1 (nothing in common). We also report the running time of the pipelines, and compare them to the running time for standard species tree estimation methods on the same datasets.

We explore GTM within the same divide-and-conquer pipeline strategy used in [3, 4], as described in Section 4, using the centroid edge decomposition and decomposing until each subset has at most 120 species.

### 5.2 External methods

We compare the trees computed using the pipelines to unpartitioned maximum likelihood using RAxML and FastTree2; we also computed trees on the multi-locus datasets using summary methods (i.e., methods that estimate the species tree by combining the gene trees) ASTRAL, NJst [13], and ASTRID [20]. NJst, ASTRID, and ASTRAL are statistically consistent under the MSC+GTR model, but FastTree2 and RAxML are not.

The divide-and-conquer pipelines require external methods to compute starting trees and constraint trees: we used FastTree2, NJst, and ASTRAL for the starting trees, and ASTRAL and RAxML for the constraint trees. We include ASTRAL and RAxML because these are the current leading methods for species tree estimation on large datasets. We include NJst and ASTRID because they, like ASTRAL, are species tree estimation methods that are polynomial time, statistically consistent under the MSC model, and can run on 1000-species datasets. Finally, we include FastTree2 (henceforth called FastTree) because it is a very fast maximum likelihood heuristic.

### 5.3 Datasets

Our study makes use of the 1000-taxon multi-locus simulated datasets from the NJMerge study [3], which are available at [21]. Here we briefly describe the experimental protocol used in [3], and how we used the datasets to test GTM pipelines.

Each replicate dataset in [3] has a true species tree and 1000 true gene trees. The gene trees were generated by evolving them down the species tree under the MSC model, which has the consequence that the true gene trees can differ from the true species tree due to incomplete lineage sorting (ILS). The sequence alignments in [3] were produced by evolving sequences down the true gene trees under the GTR model and gene trees were estimated on these alignments using FastTree [22]. The study in [3] includes model species trees with different levels of ILS and two types of genes (exons and introns) that have two different rates of evolution. Among the datasets available from [3], we selected 1000-taxon datasets from model conditions with two levels of ILS (low and very high) and both types of genes.

In order to stress test GTM, we explored conditions where estimating a reasonably accurate starting tree might be difficult. Since the number of genes impacts the accuracy of the starting tree (especially when ILS is high), we achieved this by including analyses with only 10 and 25 genes, which we selected by picking the first such genes from the data repository for the study.

We explored a range of fast methods for computing the starting tree, including FastTree (which can only be used when the number of genes is not too large), ASTRAL, and NJst [13]. The result was a set of starting trees that varied in accuracy, from very accurate to very inaccurate. In general, all starting trees on low ILS conditions were reasonably accurate, even with only 10 genes. However, with high ILS conditions and ten genes, all starting trees had low accuracy: the best starting trees had error rates in the 50-60% RF error range and the worst starting trees (computed using ASTRAL) had error rates between 64-66% (see Appendix C and Appendix D). Thus, all the starting trees for the 10-gene high ILS conditions are poor, and they provide the stress test we need to understand how GTM operates when given poor starting trees.

Overall, we considered 12 model conditions (three numbers of genes, two types of genes, and two levels of ILS), and each model condition has 20 replicates (Table 1).

**Table 1:** Dataset properties. AD values of ILS levels are the average Robinson-Foulds distances between the true species tree and the true gene trees. Gene Tree Estimation Error (GTEE) is the average RF error of the estimated gene trees.

|  |  |
| --- | --- |
| Number of species | 1000 |
| Number of sites per gene | 300-1500 |
| Number of genes | 10, 25, 1000 |
| ILS levels | Very High ILS (68-69% AD)<br>Low ILS (8-10% AD) |
| Gene types | Exons (38-64% GTEE)<br>Introns (26-51% GTEE) |

### 5.4 Experiments and computational resources

We performed three experiments. Experiment 1 determined the best way to compute starting trees and subset trees for each DTM method (NJMerge, TreeMerge, and GTM) within the divide-and-conquer pipeline, for each model condition (i.e., ILS level and number/type of genes). An important part of this experiment is that it allowed us to evaluate the impact of the starting tree on GTM, which is a matter of some interest to us. Experiment 2 compared the three DTM methods to each other with respect to accuracy and running time. Experiment 3 evaluates the impact of divide-and-conquer pipelines using GTM on RAxML and ASTRAL, the two leading species tree estimation methods, with respect to accuracy and running time.

All the analyses we performed were executed on the Campus Cluster at UIUC, which limits analyses to 4 hours; hence, any analysis that failed to complete within that time was discarded. However, for the 1000-gene analyses involving NJMerge and RAxML (which could not complete within 4 hours on the Campus Cluster), we report results given in [3] that used the Blue Waters supercomputer and allowed to run for 48 hours.

## 6 Results

Missing replicates in the figures typically reflect NJMerge (and in some cases ASTRAL and RAxML) failures to complete within the allowed running time (see Table 7 in Appendix D).

### 6.1 Experiment 1: Designing the DTM pipelines

We consider the accuracy of pipelines for each DTM method, selecting either RAxML or ASTRAL for constraint trees and NJst, ASTRAL, or FastTree for the starting tree (where they can be run within the stated time limits).

Results for TreeMerge and GTM on the high ILS datasets with 10 introns are shown in Figure 3; NJMerge is not shown for these data, as it failed for all these replicates (as well as for the analyses with 25 high ILS introns). This experiment shows that the most accurate results are obtained using NJst-ASTRAL-TreeMerge and NJst-ASTRAL-GTM (which are tied), and all other combinations have distinctly higher error. Interestingly, the next best analyses are obtained using FastTree-ASTRAL-TreeMerge and FastTree-ASTRAL-GTM (which are tied), but ASTRAL-ASTRAL-TreeMerge and ASTRAL-ASTRAL-GTM. have much higher error rates. Finally, the least accurate results are obtained using ASTRAL-RAxML-TreeMerge and ASTRAL-RAxML-GTM. Figure 3 also shows that the starting trees obtained using ASTRAL and constraint trees obtained using RAxML have very high error on these data, which explains these negative results. Results for other high ILS conditions show the same trends; see Appendix C. Overall, therefore, the most accurate analyses on high ILS datasets are obtained using NJst-ASTRAL-TreeMerge and NJst-ASTRAL-GTM.

**Figure 3:**
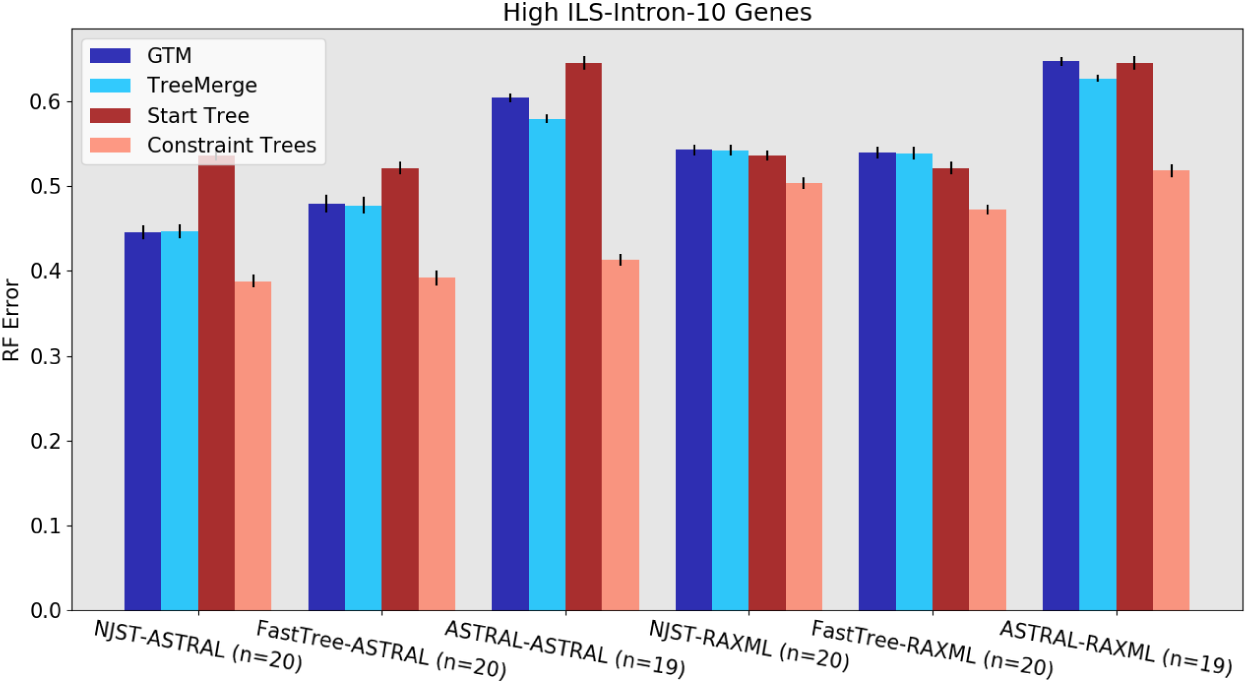
Experiment 1: Tree error rates for GTM and TreeMerge with different “starting tree-constraint tree” combinations over 10 high ILS intron genes with 1000 species. (NJMerge is not shown, because it failed for all high-ILS 10- and 25-gene replicates. Check Figure 12 for high-ILS NJMerge results). Both DTM methods work best with ASTRAL constraint trees and show a distinct preference for the NJst-ASTRAL combination. The value for *n* is the number of replicates being compared, where all methods ran. Error bars show standard error of the replicate average.

Results for low ILS datasets with 10 introns show different trends (Fig. 4). Here we see that the most accurate results are obtained using FastTree-RAxML-GTM, with FastTree-RAxML-NJMerge slightly less accurate (followed by FastTree-RAxML-TreeMerge). The next best results are obtained using other combinations with RAxML to compute the constraint trees, which is explained by noting that the RAxML constraint trees are highly accurate. We also see that the FastTree starting tree is highly accurate, which helps explain the good performance of FastTree-RAxML-GTM. Results for other low ILS conditions are shown in Appendix C, and show the same trends. Overall, for low ILS datasets, the best accuracy is obtained using FastTree-RAxML-GTM. However, FastTree cannot run to completion on 1000-gene datasets, which limits its utility as a general starting tree to only those datasets with not too many genes. Therefore, for 1000-gene datasets, we recommend NJst-RAxML-GTM.

**Figure 4:**
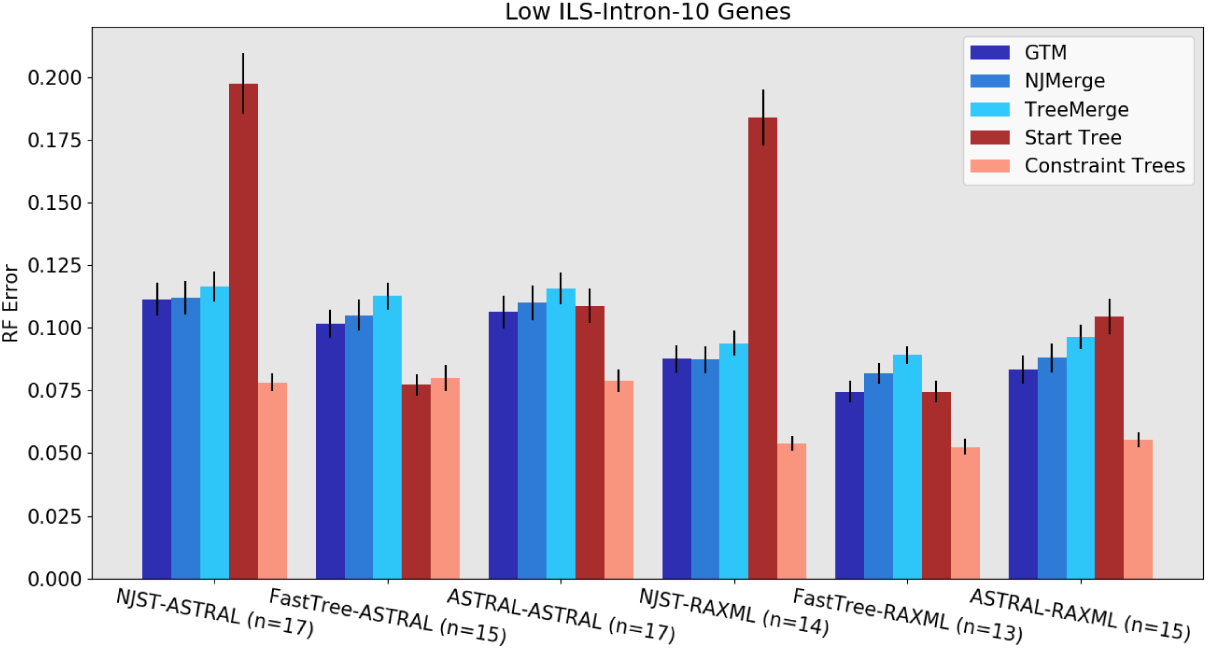
Experiment 1: Tree error rates for GTM, NJMerge, and TreeMerge with different “starting tree-constraint tree” combinations over 10 low-ILS intron genes with 1000 species. All three DTM methods work best with FastTree starting trees and RAxML constraint trees. The value for *n* is the number of replicates being compared, where all methods ran. Error bars show standard error of the replicate average.

In order for the pipeline to be provably statistically consistent under the MSC+GTR model, both the starting tree and the constraint trees must be computed using methods that are statistically consistent under the MSC+GTR model, which means (for our study) either ASTRAL or NJst for both steps. NJst is much faster than ASTRAL, and can complete on all the datasets we explored within the limited allowed time (which is not true for ASTRAL). Therefore, when statistical consistency is required, NJst-ASTRAL is the right pipeline for all DTM methods.

### 6.2 Experiment 2: Comparing DTM methods

We compare the three DTM methods on 1000-species datasets. We show results for the NJst-RAxML and NJst-ASTRAL pipelines, but results for other pipelines show similar trends.

Under the low ILS conditions, GTM and NJMerge have very similar RF error rates, with TreeMerge slightly higher in error (Fig. 5). For the high ILS conditions, the comparison is made between GTM and TreeMerge, as NJMerge failed to complete on many datasets (Table 7, Appendix D). TreeMerge and GTM have very close accuracy under most high ILS conditions, and neither dominates the other method (Fig. 16).

**Figure 5:**
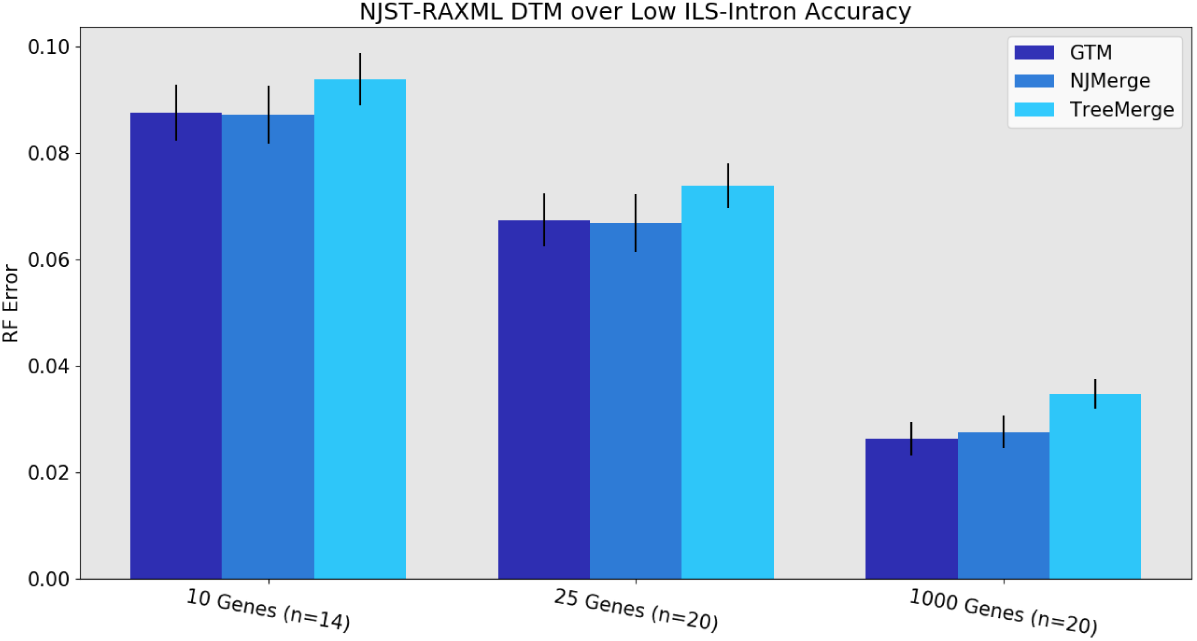
Experiment 2: Relative accuracy of different DTMs (GTM, NJMerge, and TreeMerge) within divide-and-conquer pipelines on 1000 species using NJst starting trees and RAxML constraint trees. GTM and NJMerge are about even, with a modest advantage over TreeMerge. The value for *n* indicates the number of replicates being compared, where all three methods completed. (NJMerge failed or timed out for 6 of the 10-gene replicates). Error bars show standard error of the replicate average.

**Figure 6:**
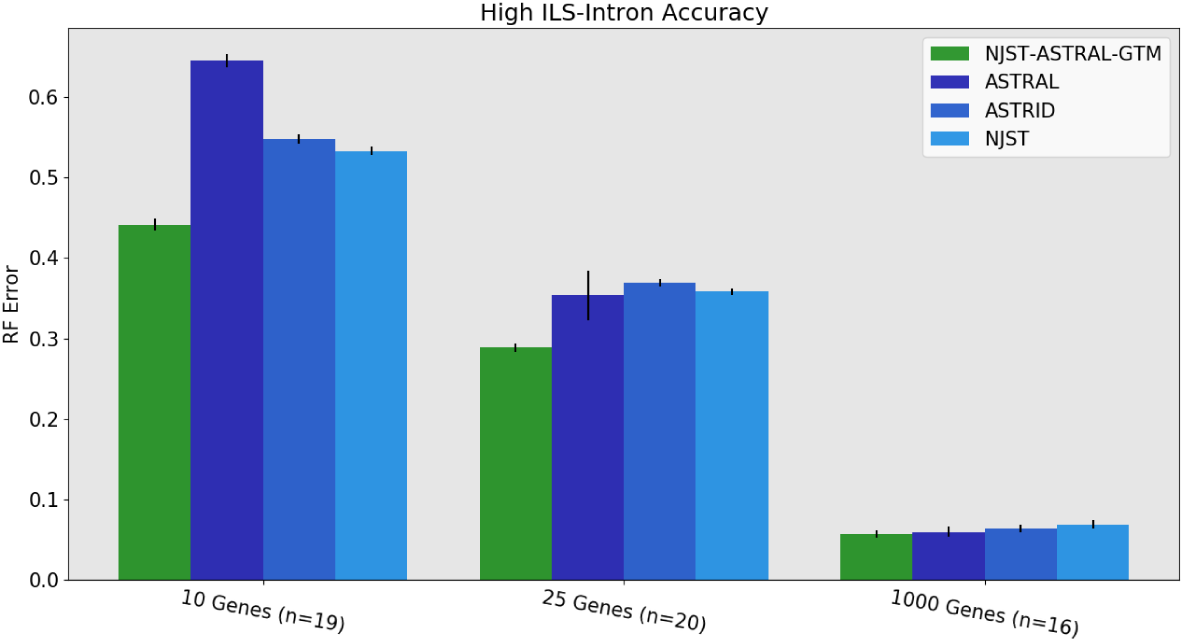
Experiment 3: Comparison of NJst-ASTRAL-GTM to AS-TRAL, ASTRID, and NJst on 1000 species with high ILS. RAxML is omitted, since it was only available for a few replicates of high ILS introns. GTM is more accurate than the other methods on small numbers of genes, and roughly matches ASTRAL and ASTRID on 1000 genes. The value for *n* indicates the number of replicates where ASTRAL trees are available. The 1000-gene ASTRAL trees were taken from [3]. Error bars indicate standard error of the replicate average.

Runtime performance is compared in Table 2 for high ILS and Table 3 for low ILS. Under conditions where NJMerge and TreeMerge both run, NJMerge is slower, and each takes many minutes (up to 30 minutes for NJMerge, and up to 10 minutes for TreeMerge). However, GTM takes under a second for all model conditions, which represents an enormous speedup over the other two methods.

**Table 2:** Average runtime (seconds) on 1000-species datasets with high ILS, using NJst starting trees and ASTRAL constraint trees. The value for *n* is the number of replicates being compared, where both GTM and TreeMerge finished. NJMerge and TreeMerge timings were unavailable for 1000 genes. NJMerge failed on all 10 and 25-gene high ILS replicates.

|  | GTM | NJMerge | TreeMerge |
| --- | --- | --- | --- |
| 10 Exons (n=20) | 0.47 | X | 655 |
| 10 Introns (n=20) | 0.55 | X | 524 |
| 25 Exons (n=20) | 0.46 | X | 504 |
| 25 Introns (n=20) | 0.46 | X | 509 |
| 1000 Exons (n=20) | 0.47 | 1950 | N/A |
| 1000 Introns (n=20) | 0.47 | 1879 | N/A |

**Table 3:** Average runtimes (in seconds) on 1000-species datasets with low ILS, using NJst starting trees and RAxML constraint trees. The value for *n* is the number of replicates being compared, where all three methods finished; NJMerge failed or timed out on the missing 10 and 25-gene low ILS replicates. TreeMerge timings were unavailable for 1000 genes.

|  | GTM | NJMerge | TreeMerge |
| --- | --- | --- | --- |
| 10 Exons (n=2) | 0.43 | 946 | 477 |
| 10 Introns (n=8) | 0.42 | 732 | 446 |
| 25 Exons (n=9) | 0.41 | 728 | 583 |
| 25 Introns (n=10) | 0.43 | 685 | 543 |
| 1000 Exons (n=20) | 0.43 | 2055 | N/A |
| 1000 Introns (n=20) | 0.43 | 2010 | N/A |

### 6.3 Experiment 3: Evaluating GTM-boosting

One way of describing these GTM pipelines is that they are designed to improve (or “boost”) the accuracy, speed, and/or scalability of a selected species tree estimation method used to compute constraint trees (which we refer to as the “base method”). Here, we explore the impact of “GTM-boosting” on ASTRAL and RAxML, two leading species tree estimation methods.

#### 6.3.1 Impact of GTM-boosting on ASTRAL

ASTRAL is the leading method for species tree estimation when gene tree heterogeneity is the result of ILS, and it is statistically consistent under the MSC. Here we compare the NJst-ASTRAL-GTM pipeline (which is statistically consistent under the MSC) to ASTRAL, and also to ASTRID and NJst, two other summary methods that are also statistically consistent under the MSC.

Results for the intron datasets with high ILS (Fig. 6) show this pipeline has the best accuracy of all methods for 10 and 25 genes (with a large improvement especially for the 10-gene datasets), and then all four methods have essentially the same error rates on 1000 genes (with a slight advantage to NJst-ASTRAL-GTM and ASTRAL). Analyses on the same datasets and including RAxML are shown in Figure 17 (Appendix C), revealing that RAxML is less accurate than the summary methods for these high ILS conditions.

Comparing the same set of summary methods under low ILS conditions (Fig. 7) shows ASTRAL and NJst-ASTRAL-GTM are essentially indistinguishable for accuracy and dominate the next best method, ASTRID, for all numbers of genes. The difference between NJst-ASTRAL-GTM and ASTRID is large for 10 genes and decreases for larger numbers of genes. Also, ASTRID clearly dominates NJst, with a large gap between the two methods for all numbers of genes.

**Figure 7:**
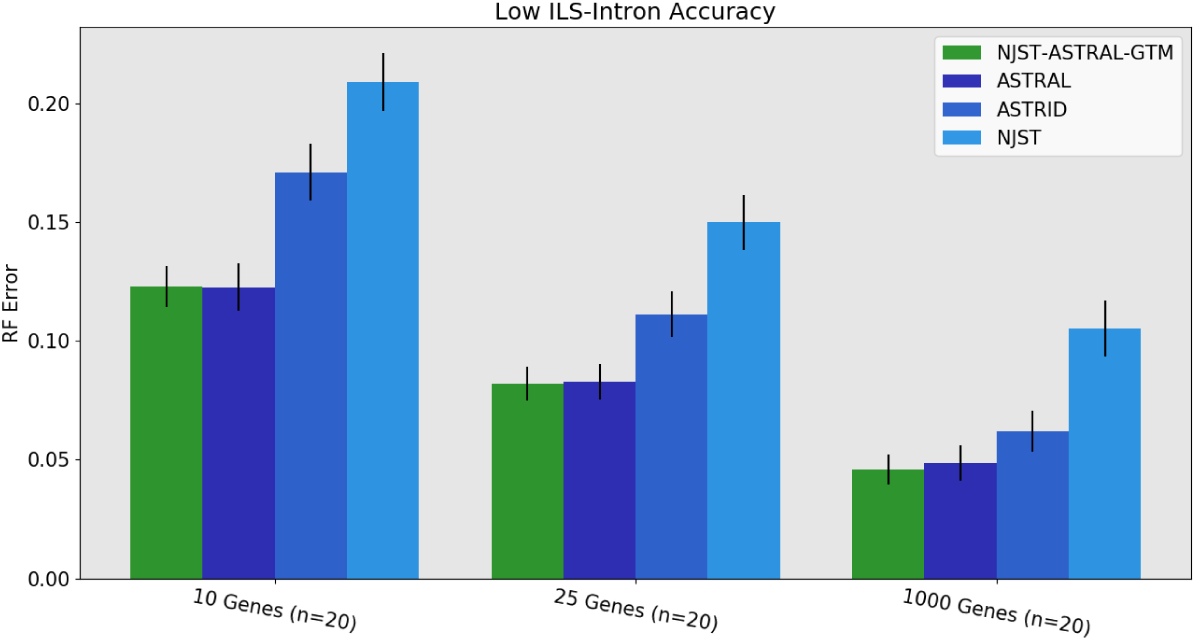
Experiment 3: Comparison of NJst-ASTRAL-GTM to AS-TRAL, ASTRID, and NJst on 1000 species with low ILS. The value for *n* indicates the number of replicates where ASTRAL trees are available. The 1000-gene ASTRAL trees were taken from [3]. Error bars indicate standard error of the replicate average.

Next, we compare the running time for NJst-ASTRAL-GTM and ASTRAL. Under high ILS conditions (see Table 4), ASTRAL took 2.4 hours on 10 introns whereas NJst-ASTRAL-GTM completed in 98 seconds, so that ASTRAL uses almost two orders of magnitude more time than NJst-ASTRAL-GTM. The magnitude of the reduction in running time decreased as the number of genes increased, but was still large for 1000 genes (42.5 hours, for ASTRAL and 2.2 hours for NJst-ASTRAL-GTM). These are dramatic savings in running time.

**Table 4:**
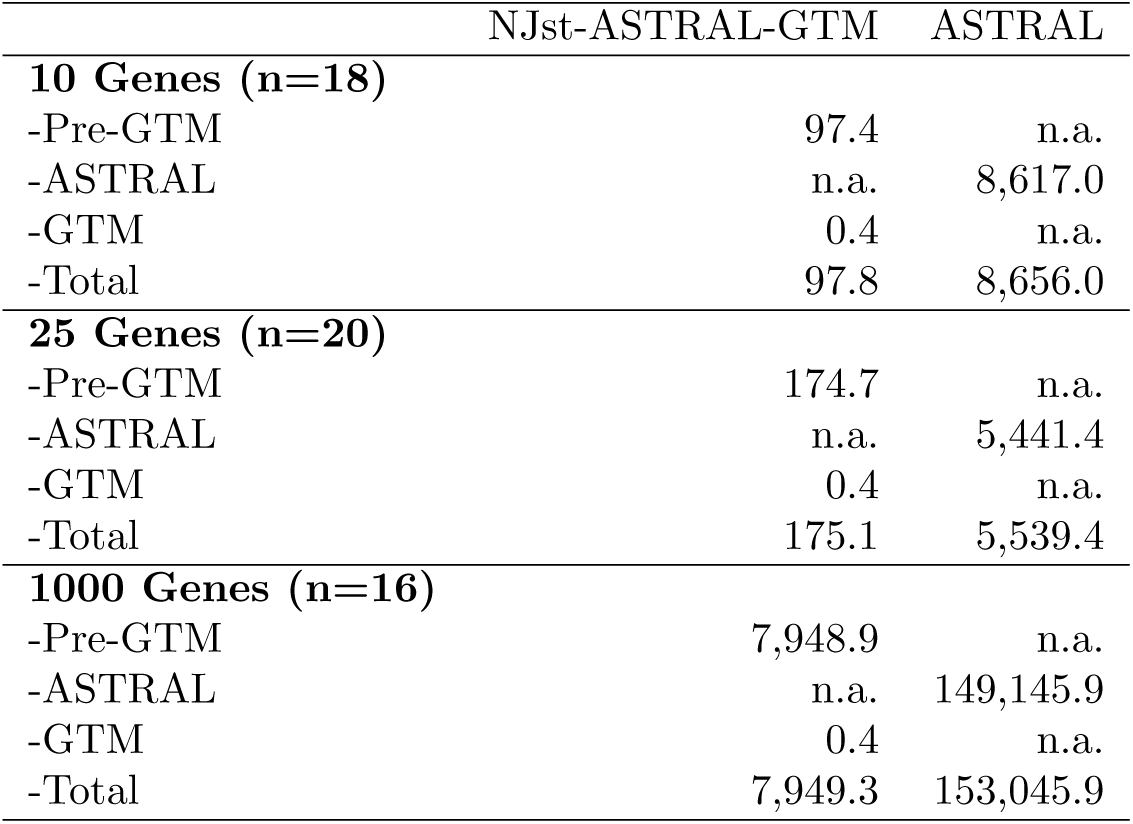
Comparison of average runtime (seconds) of NJst-ASTRAL-GTM and ASTRAL for high ILS conditions with introns on 1000 species. The value for *n* is the number of replicates being compared (i.e., where ASTRAL trees are available). Pre-GTM covers computing gene trees using FastTree, the NJst starting tree, and ASTRAL subset trees; the gap between “total” and “ASTRAL” for the right hand column reflects the time to compute gene trees using FastTree, which is 3.9 seconds per gene. Results for the 1000-gene ASTRAL trees are taken from the NJMerge study [3].

The difference is less dramatic under low ILS conditions, since ASTRAL runs much faster. Nevertheless, the pipeline runs a bit more than twice as fast on 10 genes, a bit less than twice as fast on 25, and about twice as fast on 1000 (Table 5).

**Table 5:** Comparison of average runtime (seconds) of NJst-ASTRAL-GTM and ASTRAL for low ILS conditions with introns on 1000 species. The value for *n* is the number of replicates being compared (i.e., where ASTRAL trees are available). Pre-GTM covers computing gene trees using FastTree, the NJst starting tree, and ASTRAL subset trees; the gap between “total” and “ASTRAL” for the right hand column reflects the time to compute gene trees using FastTree, which is 3.9 seconds per gene. Results for the 1000-gene ASTRAL trees are taken from the NJMerge study [3].

|  | NJst-ASTRAL-GTM | ASTRAL |
| --- | --- | --- |
| **10 Genes (n=20)** |  |  |
| -Pre-GTM | 59.1 | n.a. |
| -ASTRAL | n.a. | 114.4 |
| -GTM | 0.4 | n.a. |
| -Total | 59.5 | 153.4 |
| **25 Genes (n=20)** |  |  |
| -Pre-GTM | 120.9 | n.a. |
| -ASTRAL | n.a. | 101.3 |
| -GTM | 0.4 | n.a. |
| -Total | 121.3 | 199.3 |
| **1000 Genes (n=19)** |  |  |
| -Pre-GTM | 4,308.4 | n.a. |
| -ASTRAL | n.a. | 4,894.4 |
| -GTM | 0.4 | n.a. |
| -Total | 4,308.8 | 8,794.4 |

**Table 6:**
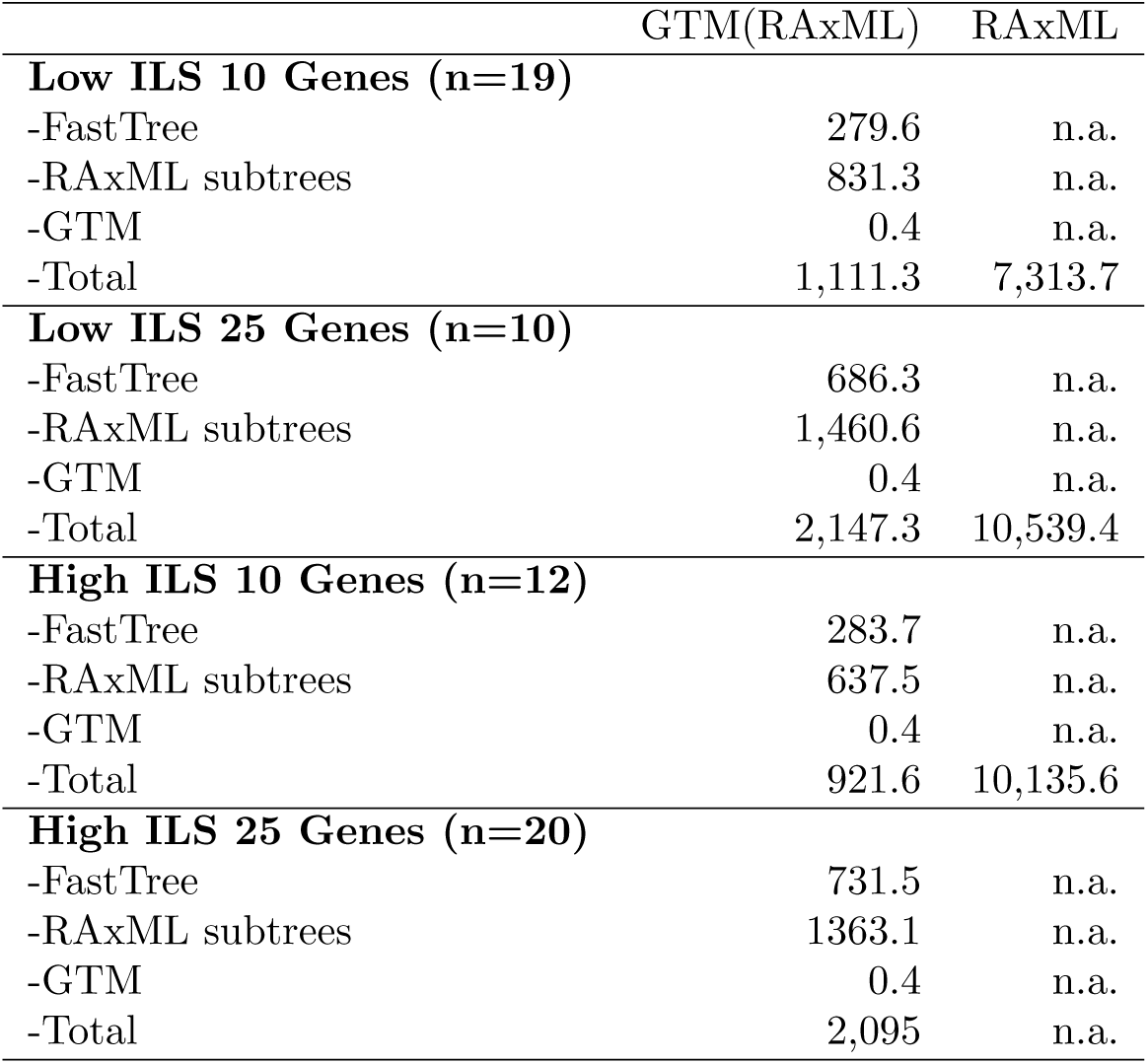
Average runtime (seconds) of FastTree-RAxML-GTM (GTM(RAxML)) and RAxML on 1000-species exon datasets. The value for *n* is the number of replicates being compared, i.e., where a RAxML tree is available.

#### 6.3.2 Impact of GTM-boosting on RAxML

Concatenated analysis using maximum likelihood is one of the major approaches to species tree estimation. RAxML is considered by many to be the leading ML heuristic, and outperforms FastTree, a fast but less accurate maximum likelihood heuristic, with respect to maximum likelihood scores. However, FastTree can run on very large datasets, and can (in some conditions) match the topological accuracy of RAxML [23].

Here we address the potential for scaling RAxML to large datasets through the use of a divide-and-conquer pipeline using GTM. As we noted earlier, FastTree-RAxML-GTM has higher accuracy than NJst-RAxML-GTM but cannot complete (within the 4 hour time limit) on the 1000-gene datasets, and for this reason we also include NJst-RAxML-GTM. We compare these two versions of GTM-boosting to RAxML and FastTree, under both low and high ILS conditions.

When restricted to the datasets where FastTree completes, we see the following trends. Under low ILS, FastTree-RAxML-GTM provides substantially better accuracy than NJst-RAxML-GTM and is nearly identical to RAxML, although it is slightly worse than FastTree by itself (Fig. 8). Under high ILS, FastTree-RAxML-GTM and NJst-RAxML-GTM are indistinguishable for accuracy, and both are somewhat more accurate than RAxML and less accurate than FastTree (Fig. 9). The very good accuracy of FastTree in comparison to RAxML is noteworthy, especially since RAxML is established to be a better maximum likelihood heuristic than FastTree. However, due to computational limitations in this study, we ran RAxML with only one random starting condition, instead of running it with a larger number (e.g., 10 or 20) of random starting conditions and then selecting the tree with the best ML score. This choice may have reduced the accuracy of RAxML relative to FastTree.

**Figure 8:**
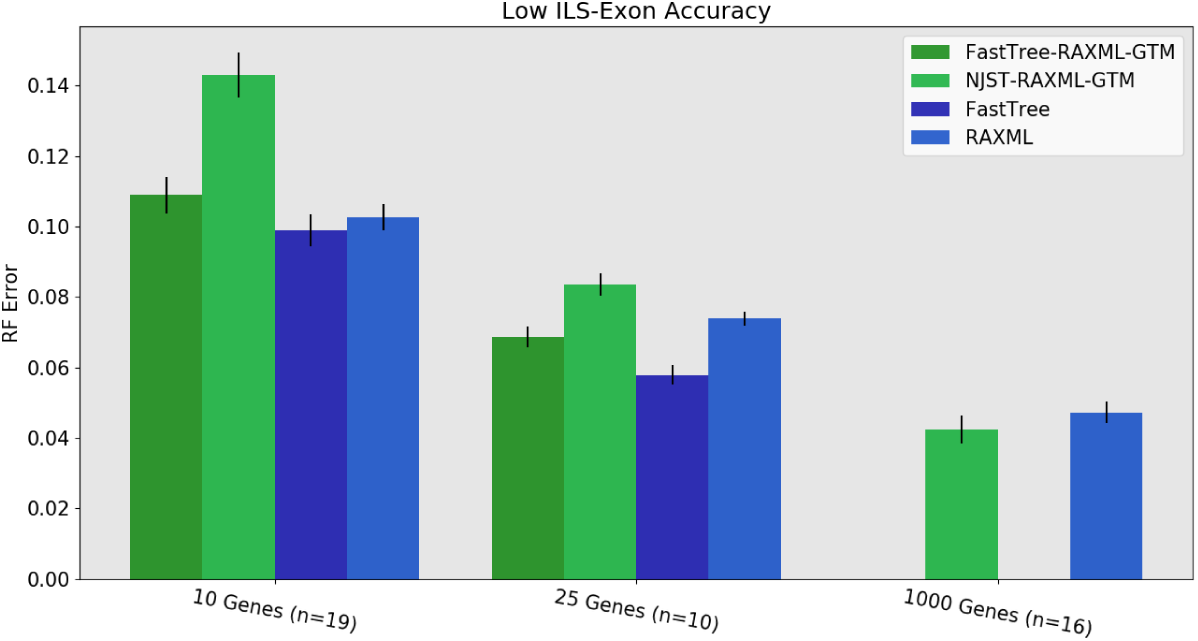
Experiment 3: Comparison of FastTree-RAxML-GTM and NJst-RAxML-GTM to RAxML and FastTree on 1000-species datasets with low ILS exons. The value for *n* is the number of replicates on which RAxML completed; missing replicates indicate RAxML exceeding run-time limits on 10 and 25 genes (the 1000-gene RAxML trees are taken from [3]). FastTree was not used for 1000 genes. Error bars show standard error of the replicate average.

**Figure 9:**
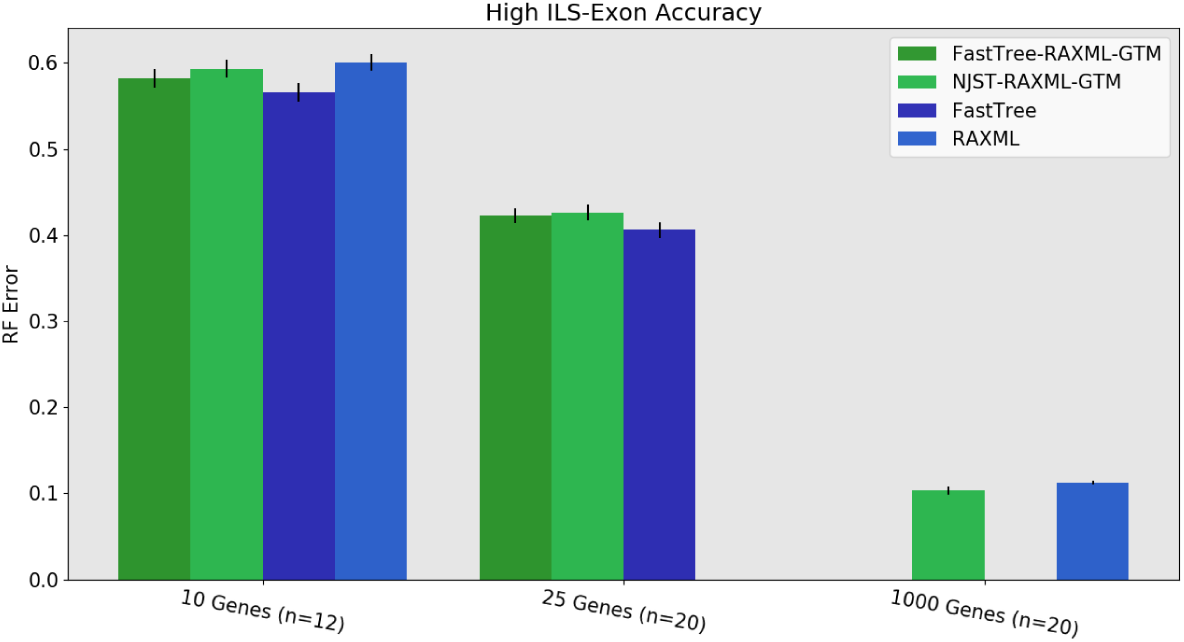
Experiment 3: Comparison of FastTree-RAxML-GTM and NJst-RAxML-GTM to RAxML and FastTree on 1000-species datasets with high ILS exons. The value for *n* is the number of replicates on which RAxML completed; missing replicates indicate RAxML exceeding run-time limits on 10 and 25 genes (the 1000-gene RAxML trees are taken from [3]). FastTree was not used for 1000 genes. Error bars show standard error of the replicate average.

We now discuss the running time comparisons between RAxML and the two GTM pipelines that use RAxML to compute constraint trees. RAxML often failed to complete within the allowed time limit (4 hours) on many 10- and 25-gene datasets. Results reported on the 1000-gene datasets were obtained in [3], where RAxML was allowed to run for 48 hours, and the best found tree was returned. However, even on the 10-gene and 25-gene datasets there are noteworthy differences, which we now discuss. FastTree-RAxML-GTM and NJst-RAxML-GTM completed on all the 10- and 25-gene datasets but RAxML completed on only 15/80 25-gene datasets and 55/80 10-gene datasets (Table 7 in Appendix D).

When restricted to those datasets where RAxML completed within 4 hours, RAxML used (on average) between 2 and 3 hours, depending on the model condition (with more time needed for the high ILS datasets with 25 genes). In comparison, the average running time on these datasets for the two GTM pipelines were much smaller: at most 27 minutes for NJst-RAxML-GTM and at most 36 minutes for FastTree-RAxML-GTM. Thus, NJst-RAxML-GTM is the fastest, followed (fairly closely) by FastTree-RAxML-GTM, and both are much faster than RAxML. However, because FastTree cannot run on the 1000-gene datasets, NJst-RAxML-GTM is the only scalable GTM pipeline that uses RAxML on subsets.

## 7 Discussion

The datasets generated for this study each have 1000 species but otherwise vary in terms of number of genes (10, 25, and 1000), ILS level (low and high), and type of gene (exon or intron). Thus, the model conditions provide a range of conditions that include many of the conditions observed in species tree estimation for large numbers of species. That said, the observations made in this study are limited to these conditions, and other conditions may show other trends.

The first overall observation we make is that GTM is comparable in accuracy to TreeMerge and NJMerge (but is much faster). We expected to see an improvement in running time over NJMerge, which is not designed to be very fast, but we did not expect to necessarily see an improvement in running time over TreeMerge, and the degree of improvement was larger than we might have expected. The observation that GTM is typically comparable in accuracy to TreeMerge and NJMerge is surprising because GTM cannot blend, while TreeMerge and NJMerge can.

We focus on the comparison between TreeMerge and GTM, since NJMerge failed on many datasets in our experiment. There are some conditions where GTM is slightly more accurate than TreeMerge and some other conditions where GTM is slightly less accurate than TreeMerge, and these conditions have different properties. Specifically, there can be a small advantage to GTM when the starting tree is not too poor (at most 60% RF error) and a small advantage to TreeMerge when the starting tree has very poor accuracy (i.e., on the 10-gene high ILS datasets, where the ASTRAL starting trees had 64-65% RF error rates, see Figs. 10 and 3). Thus, the accuracy of the starting tree has an impact on the relative accuracy of GTM and TreeMerge. We expected to see GTM degrade in accuracy with poor starting trees (which is why we included conditions where poor starting trees were computed), and so this part is not surprising. However, what *is* surprising is that GTM matches the accuracy of TreeMerge even with fairly mediocre starting trees, including ones where the starting tree has 50-60% RF error. Overall, our study shows that the limitation of not being able to blend subset trees was not a significant problem for GTM, under the conditions we explored.

The next observations have to do with the impact of “GTM-boosting” on the method used to compute constraint trees (i.e., the “base method”). Our study shows that GTM-boosting provides substantial improvements in speed for all tested model conditions and generally maintained (and sometimes even improved) accuracy over the base method. The improvement in running time obtained by GTM-boosting was expected, as previous divide-and-conquer approaches have shown similar trends [24, 25].

The improvement in accuracy seen in NJst-ASTRAL-GTM over ASTRAL and NJst-RAxML-GTM or FastTree-RAxML-GTM over RAxML are surprising, and worth trying to understand.

The conditions in which NJst-RAxML-GTM or FastTree-RAxML-GTM were more accurate than RAxML were cases with 25 genes, where they produced very slightly better results; for other numbers of genes, RAxML was more accurate. Somewhat similar trends were observed in [3] when NJMerge was used with RAxML in the same pipeline as we used here and found to produce somewhat more accurate trees than RAxML. The explanation offered in [3] was that NJmerge+RAxML is a combination of a coalescent-based species tree estimation method (the starting tree) and concatenation analysis method (the constraint trees), and the combination allows it to be more accurate than a method that is purely concatenation-based. In our study, this advantage does not hold for the smaller numbers of genes, where NJst-RAxML-GTM is much less accurate than RAxML for 10 genes under the low ILS condition. Our explanation for why NJst-RAxML-GTM is substantially less accurate than RAxML for 10-gene low ILS datasets is that the NJst starting tree has high error (Fig. 13, Appendix C). This is consistent with the observations that FastTree-RAxML-GTM is close to RAxML for accuracy on low ILS conditions, even with 10 genes, and the FastTree starting tree also has high accuracy. Thus, as we have seen, the starting tree has an impact on accuracy, and the choice of starting tree should reflect the level of ILS.

The conditions in which NJst-ASTRAL-GTM was more accurate than ASTRAL occur for the high ILS model conditions with ten genes. Frankly, this improvement in accuracy is surprising and was not expected. However, Figures 3 and 10 (Appendix C) showing results for high ILS datasets with 10 introns and 10 exons, respectively, provide some insight. The difference in tree error for ASTRAL on the full dataset (i.e., as the starting tree) and on the subsets (i.e., as the constraint tree method) are dramatic for both introns and exons. For example, ASTRAL had average 64% RF error on 10 introns but the average ASTRAL constraint tree error was only 39% error—a large drop in error rate. In other words, ASTRAL was more accurate on the smaller subsets (each with at most 120 species) than it was with the full dataset (with 1000 species). One possible explanation for this trend is that when the number of genes is very small, the constrained search strategy employed by ASTRAL works better with a smaller number of species.

GTM using appropriate constraint tree methods (i.e., ASTRAL for high ILS and RAxML for low ILS) nearly always improved on the starting tree, often substantially. The only exceptions to this rule are the low ILS conditions with 10 or 25 genes where the best starting trees had very low error (Fig. 4, and also Figs. 1 0-15, Appendix C). Even for these cases, using GTM matched the starting tree accuracy except for one model condition (25 low ILS exon datasets) where it produced a tree with 1% higher error (Fig. 15). In contrast, the other DTMs reduced accuracy compared to these highly accurate starting trees.

Our study also showed that GTM divide-and-conquer pipelines reduce the running time and enable expensive base methods (here, RAxML and ASTRAL) to be applied to large datasets. The improvement in running time is most noticeable for the high ILS conditions, where ASTRAL and RAxML require more time. Furthermore, for all numbers of genes and ILS conditions, the vast majority of the time used within a GTM pipeline is the pre-GTM component (which computes the gene trees, the starting tree, and the subset trees); in contrast, the GTM part used approximately half a second, even on 1000 genes and 1000 species.

A basic question that is relevant to GTM divide-and-conquer pipelines is the sensitivity of the approach to the algorithmic parameters, including the starting tree (which is also used as the guide tree), the method used to compute constraint trees, and even the decomposition strategy. Our study shows that the starting tree and the constraint tree method both have an impact on the accuracy of the final tree. For example, we noted that FastTree-RAxML-GTM is more accurate than NJst-RAxML-GTM on low ILS datasets (reflecting the improved accuracy of FastTree for low ILS conditions), and that NJst-ASTRAL-GTM is more accurate than NJst-RAxML-GTM on high ILS datasets (reflecting the improved accuracy of ASTRAL as a constraint tree method for high ILS conditions). However, we did not evaluate changes to the decomposition strategy (e.g., changing the subset size or which edges we delete) nor did we examine the potential impact of allowing the guide tree to be different from the starting tree.

One obvious case where the starting tree and guide tree could be different is where GTM is used within an iterative strategy, where each iteration uses the tree computed during the previous iteration for the starting tree, divides into subsets, constructs trees on the subsets, and then merges the constraint trees using a fixed guide tree. In such a scenario, the guide tree is fixed but the “starting tree” (which is only used to compute the decomposition) can change in each iteration. In this context the guide tree only impacts the merger step and not also the decomposition strategy, and the impact of the guide tree (when it is used with a good starting tree) is an interesting question. For example, if the decomposition is based on the true tree and the constraint trees are the true tree on the subsets, then how much can a bad guide tree impact the outcome? A simple thought experiment, based on only dividing into two subsets, reveals that the impact may not be minor. For example, consider the case where the true tree is a caterpillar tree with *n* leaves (i.e., a path of length *n* − 3 with the leaves hanging off the path) and the division is into two subsets. Given a random guide tree, GTM would combine the two trees by subdividing a random edge in each of the two constraint trees (thus creating two nodes, *v*_1_ and *v*_2_), and attaching the two trees by adding a new edge between *v*_1_ and *v*_2_. Note that such a randomly merged tree would have high error, even though the constraint trees and the starting tree are completely accurate. Thus, a random guide tree can result in high error, though the degree of error will depend on the topology of the true tree, the decomposition strategy (and in particular the number of subsets), and the error in the constraint trees.

## 8 Summary and Conclusions

Divide-and-conquer phylogeny estimation using Disjoint Tree Merger (DTM) is a recently developed approach to large-scale tree estimation that has been shown to improve speed while maintaining (and in some cases improving) accuracy for large-scale phylogenomic datasets, and maintaining statistical consistency under the MSC+GTR model. Here, we proposed the DTM-GT problem, which seeks a compatibility tree that is as close as possible to an input “guide tree”. We proved that DTM-GT is NP-hard, and presented GTM, which solves the problem exactly and in polynomial time if blending is not allowed. Our experimental study showed that GTM generally matches or improves on the accuracy of two prior DTM methods, TreeMerge and NJMerge, while being much faster. Therefore, although GTM has the vulnerability of being unable to blend (a limitation that neither TreeMerge nor NJMerge have), this vulnerability had limited impact in this study.

Importantly, GTM-boosting improved the accuracy and speed of ASTRAL, a leading method for species tree estimation that addresses gene tree heterogeneity due to incomplete lineage sorting. GTM-boosting also improved the speed for RAxML, the leading heuristic for concatenated maximum likelihood, while coming very close to it in accuracy (and sometimes being even more accurate) when used with FastTree as the starting tree. Thus, GTM-boosting provides substantial advances for multi-locus species tree estimation on large numbers of species.

Our study reveals the surprising potential of unblended DTM methods, and suggests many directions for future research. For example, GTM is just the first unblended DTM, and it seems likely that other unblended DTMs might be more accurate. Since GTM is impacted by its guide tree, unblended DTMs that are not based on guide trees might be more accurate.

GTM itself could be modified to allow for blended merging, with potential improvement in accuracy. For example, the refinement step in GTM where the collapsed tree is refined to separate the constraint sets is very simple, and is performed to ensure an unblended merger. However, if blending were permitted, then other ways of refining the tree could be considered (provided that they do not produce trees that violate the constraint trees). Polytomy refinement is a natural problem in phylogenetics (e.g., [26]), and has also been specifically discussed in the context of gene tree correction given a species tree; see, for example, [27] (and references therein) for a discussion of refinement techniques that address ILS and [28] (and references therein) for polytomy refinement techniques that address gene duplication and loss.

DTM methods (blended or unblended) can also be used for other phylogeny estimation problems where the most accurate base methods are computationally intensive. For example, DTMs could also be used with Bayesian methods to co-estimate gene trees and species trees [29] or for “genome rearrangement phylogeny”, which takes chromosomal rearrangements, duplications, and other events that change the chromosomal architecture into account.

Furthermore, a parallel implementation of the divide-and-conquer pipeline could have high impact. As our study showed, the most expensive part of these divide-and-conquer pipelines is the computation of the constraint trees. With sufficient parallelism, these analyses could become very fast, and each iteration could complete quickly, leading potentially to the ability to compute species trees on ultra-large datasets (with thousands of genes and species) within a few hours.

Finally, we note that there are cases where an adequately accurate starting tree cannot be estimated using currently available fast methods. Examples of such situations include the challenge of estimating a tree from ultra-large datasets of unaligned sequences where alignment estimation is difficult due to sequence heterogeneity and size; this challenge is exacerbated when the datasets include fragmentary sequences, which are not only difficult to align but can reduce accuracy for tree estimation [30, 31]. For such data, standard two-phase methods that first compute an alignment and then compute a tree do not have acceptable accuracy, while PASTA [32], BAli-Phy [33], and other co-estimation methods are not fast. It is possible that alignment-free methods (see [34, 35, 36] for an entry into this topic) might provide good starting trees, but these have not been tested on ultra-large datasets (with thousands of species), and have instead mainly been focused on genome-scale analyses of tens of genomes. However, for any large dataset on which the starting trees cannot be reasonably accurately estimated quickly, blended DTM divide-and-conquer strategies may provide the best accuracy. Thus, future work into developing new DTMs, both blended or unblended, is merited.

We close with some comments about the implications of this study for largescale phylogenomic analysis. As shown in this study, the choice of method for species tree estimation depends on the level of ILS and even the number of genes. The results for low ILS datasets suggest that maximum likelihood heuristics, such as FastTree and RAxML, or using GTM-boosting with fast ML heuristics for the starting tree and more accurate ML heuristics for constraint trees, may be advantageous. Of interest is the relative performance of RAxML and FastTree (which favored FastTree in our experiments), but we note that RAxML was not run so as to obtain the best accuracy, and so further study is needed. Results for high ILS datasets are more definitive: GTM-boosting, using NJst as the starting tree and ASTRAL for constraint trees, matched or improved on ASTRAL for all model conditions and numbers of genes, and was always faster. Thus, for high ILS conditions, GTM-boosting provides a distinct advantage for both accuracy and time.

## Declarations

### Availability of data and material

Datasets used can be found at https://databank.illinois.edu/datasets/IDB-3204101 and https://databank.illinois.edu/datasets/IDB-1424746. GTM software available at https://github.com/vlasmirnov/GTM.

### Authors’ contributions

TW directed the research. VL designed and implemented the algorithm, performed the experimental study, and created the figures. VL and TW proved the theorems and wrote the paper.

#### Acknowledgements

The authors thank Sarah Christensen, Erin Molloy, Pranjal Vachaspati, and Xilin Yu for helpful comments.

## Authors’ information

VL email is and TW email is.

## A Codes used in this study, with commands

### A.1 Centroid edge decomposition strategy

We use the centroid edge decomposition strategy to divide our species sets into disjoint subsets; this is the same decomposition strategy used in the NJMerge [2, 3] and TreeMerge [4] studies. The centroid edge decomposition technique originated in SATé-II [37] and has also been used in PASTA [38, 32] and DACTAL [39]. The basic objective of this strategy is to produce a decomposition where every subset is “local” within the input tree, and does not exceed a specified maximum subset size.

The centroid edge decomposition is based on finding and deleting centroid edges, which are edges whose removal splits the leaf set into two sets where the difference in size is minimized among all the edges in the tree; because there can be more than one edge with this property, the centroid edge may not be unique.

In the centroid edge decomposition strategy, a tree *T* is given as well as a bound *B* on the size of the subsets to be computed. If *T* has at most *B* leaves, then the set of leaves of *T* is returned (i.e., there is no decomposition performed). Otherwise, a centroid edge is found and removed, and then the algorithm recurses on the two subtrees. At the end of this recursive strategy, the set of leaves of *T* (i.e., taxa) is decomposed into disjoint sets, each of which has size at most *B*.

### A.2 Codes for computing trees

#### ASTRAL 5.6.3

Code available at https://github.com/smirarab/ASTRAL

Command:

~~~
java -jar astral.5.6.3.jar
-i gene_trees.tre -o output_tree.tre
~~~

#### ASTRID 1.4

Code available at https://github.com/pranjalv123/ASTRID-1

Command:

~~~
ASTRID-linux -i gene_trees.tre
-o output_tree.tre -c output_matrix.mat
~~~

#### FastTree 2.1

We ran FastTree in an unpartitioned analysis (i.e., treating all the loci as evolving down the same model tree).

Command:

~~~
FastTree -nt -gtr
alignment.fas > output_tree.tre
~~~

#### NJMerge

Code available at https://github.com/ekmolloy/njmerge

Command:

~~~
python3 njmerge.py
-t constraint_tree1.tre constraint_tree2.tre
-m distance_matrix.mat -o output_tree.tre
~~~

#### NJst

To compute the NJst tree, we use ASTRID (with the “-c” flag) to compute the matrix of average (across all gene trees) topological distances between species. We then compute a neighbor joining [12] tree using FastME v. 2.15 [40] with command:

~~~
fastMEPath -mN
-i distance_matrix.mat -o output_tree.tre
~~~

#### RAxML 8.2.10

We ran RAxML in an unpartitioned analysis (i.e., treating all the loci as evolving down the same model tree), and with only one random starting condition.

Command:

~~~
raxmlHPC-PTHREADS-SSE3
-m GTRGAMMA -F -p 12345
-n output -s alignment.fas -T 4
~~~

#### TreeMerge

Code available at https://github.com/ekmolloy/treemerge

Command:

~~~
python3 treemerge.py
-s start_tree.tre
-t constraint_tree1.tre constraint_tree2.tre
-m distance_matrix.mat -o output_tree.tre
-p paup4a165_centos64 -w workingDir
~~~

### A.3 Other details

#### A.3.1 Obtaining a decomposition (1 round)

We used the script build subsets from tree.py, found in tools.zip in the NJMerge data repository at https://databank.illinois.edu/datasets/IDB-1424746. The method was executed by calling the decompose trees function in the script with the tree object to decompose and a maximum subset size (120). However, the script can also be run through its main method by specifying the maximum subset size and the input tree file.

#### A.3.2 Applying ASTRAL to subsets of species

To construct ASTRAL trees on subsets, we first restrict each of the gene trees to the specified subset, then we run ASTRAL on the set of restricted gene trees. To restrict each gene tree to the relevant subset, we use the Dendropy [41] command `retain_taxa_with_labels(<subset taxa>)`. The resulting restricted gene trees are written file using Dendropy’s as string(schema=“newick”) function, and ASTRAL is called with that file as input.

#### A.3.3 Computing RF distance

To compute the RF distance between two trees, each on *N* leaves, we ran the compare trees.py script, found in tools.zip in the NJMerge data repository at https://databank.illinois.edu/datasets/IDB-1424746, using the following command:

~~~
compare_trees.py <tree1> <tree2>
~~~

To return the RF error rate, we divided these values by 2*N* − 6.

## B Additional proofs

### B.1 Proof of Theorem 1

**Theorem 1** DTM-GT-FN is NP-hard

*Proof.* We first define the decision problem, **DTM-GT-FN-0**: given the same inputs as DTM-GT-FN, is there a tree *T* ′ with 0 FN distance from the guide tree *T* *? We will now show that DTM-GT-FN-0 is NP-hard, from which the NP-hardness of DTM-GT-FN naturally follows.

We reduce the Unrooted Tree Compatibility problem to DTM-GT-FN-0. The Unrooted Tree Compatibility decision problem is given a set of unrooted trees on overlapping leaf sets, determine if there exists a tree that induces them all. Unrooted Tree Compatibility is known to be NP-complete [42] (see also discussion in [43]). so this reduction will establish the NP-hardness of DTM-GT-FN-0. Let *T*_1_, *T*_2_, … *T*_*k*_ be unrooted binary trees on overlapping subsets of leaf set *L*. The following polynomial-time procedure will convert these inputs to an instance of DTM-GT-FN-0.

- First, we define a new leaf set *L*′ as follows: for each leaf *l* ∈ *L* that appears in n trees 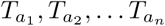 will contain n leaves 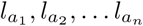.
- Second, we produce a set of disjoint constraint trees on *L*′: for each tree *T*_*i*_, we produce a constraint tree 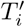 by replacing each leaf *l* ∈ *L* in *T*_*i*_ with leaf *l*_*i*_ ∈ *L*′. Since each leaf in each constraint tree is indexed by the tree it’s in, the trees are trivially disjoint.
- Third, we produce our guide tree *T* *: we include the trivial bipartition for each leaf in *L*′, and, for each leaf *l* ∈ *L* that corresponds to 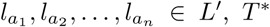 gets one bipartition separating 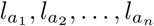 from all other leaves: 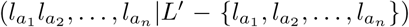. Topologically, we can build such a tree by starting with a star tree on the leaves in *L*, and replacing each leaf *l* with an internal node, whose children are the corresponding leaves 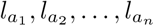. Note that the construction of the constraint trees and the guide tree takes polynomial time and their size is bounded by a polynomial in the input to Unrooted Tree Compatibility. To complete the proof, we need to verify that solving DTM-GT-FN-0 on our guide tree *T* * and our new constraint trees 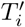 yields “yes” if and only if Unrooted Tree Compatibility yields “yes” on *T*_*i*_.

If there exists some supertree *S* that induces all trees *T*_*i*_, then there exists a tree *R* with 0 FN distance from *T* *: we obtain *R* from *S* in the same fashion as we produced *T* * earlier–by replacing each leaf *l* ∈ *L* in *S* with one internal node whose children are the corresponding 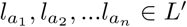. Trivially, *R* would contain all bipartitions from *T* *, and, since *S* induces each *T*_*i*_, *R* would likewise induce 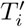. On the other hand, if such an *S* doesn’t exist, then there is no *R* with 0 FN distance from *T* *: if it did, we could recover *S* from *R* by performing the opposite operation.

Thus, the Unrooted Tree Compatibility problem reduces to DTM-GT-FN-0, showing that DTM-GT-FN-0 is NP-hard, and, therefore, DTM-GT-FN is NP-hard.

□

### B.2 Proof of Theorem 2

**Theorem 2** Given a guide tree *T* * and set 𝒯 = {*T*_1_, *T*_2_, …, *T*_*k*_} of constraint trees over their respective leaf sets *L*(*T*_*i*_) of 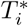, GTM optimally solves Unblended DTM-GT-FN on (𝒯, *T* *) and has a worst case running time of *O*(*N* ^2^) where *N* = |*L*(*T* *)| is the total number of species.

*Proof.* We begin with proving correctness (i.e., that GTM finds an optimal solution to DTM-GT-FN), and then prove the running time.

(1) Proof of correctness: It is clear by construction that GTM produces an unblended compatibility supertree for 𝒯. The only issue therefore is whether it returns a tree that has the smallest FN rate of all such supertrees. Then, given the recursive structure described above, it will suffice to show that each output of *Rejoin*(*T*) satisfies all constraint trees *T*_*i*_ such that *L*(*T*_*i*_) ⊆ *L*(*T*) and contains all bipartitions in *T*. Recall that we collapsed all violating edges in *T* *, as well as any edges that violate convexity. These two types of edges obviously may not appear in any valid “unblended joining” result, so if *Rejoin*(*T*) contains all bipartitions of *T*, it achieves the best possible FN distance to *T* *|*L*(*T*).

The base case is when *L*(*T*) corresponds to some *L*(*T*_*i*_), and *Rejoin*(*T*) just returns the constraint tree, *T*_*i*_. This case is trivially correct. Then, we can look at the inductive step, which is as follows. We divide *T* into *S*_1_ and *S*_2_ on bridge edge *e*. Our inductive hypothesis is that 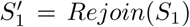 and 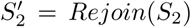 satisfy all constraint trees on their respective leaf sets and contain all bipartitions of *S*_1_ and *S*_2_. We find attachment edges 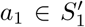 and 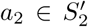 from *e*_1_ ∈ *S*_1_ and *e*_2_ ∈ *S*_2_ as described in the procedure, join them with edge *e*′ to form *T* ′, and want to prove that *T* ′ satisfies all constraint trees *T*_*i*_: *L*(*T*_*i*_) ⊆ *L*(*T*) and contains all bipartitions of *T*.

We first notice that an attachment edge *a*_1_ ∈ *S*_1_, such that *b*(*a*_1_) = *b*(*e*_1_), must always exist: since *e*_1_ ∈ *S*_1_, our inductive hypothesis says that *b*(*e*_1_) must appear in 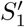. The same goes for *a*_2_. Since 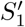 and 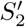 satisfy all constraint trees on their respective leaf sets and are connected by a single edge without modification to either one, then *T* ′ also straightforwardly satisfies both sets of constraint trees. Now, let *s* be any edge in *T*. We want to show that *b*(*s*) must appear in *T* ′. There are three possibilities:

1) *s* is an edge in *S*_1_.

2) *s* is an edge in *S*_2_.

3) *s* is *e*.

The third case is clear, since *e* and *e*′ are the same bipartition *L*(*S*_1_)|*L*(*S*_2_). WLOG, let’s assume the first case. This means that *b*(*s*) is a bipartition of the form *A*|(*B* ∪ *L*(*S*_2_)), where *A* ∪ *B* = *L*(*S*_1_). Then, bipartition *A*|*B* appears in *S*_1_ and, by our inductive hypothesis, there’s an edge 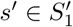, such that *b*(*s*′) = *A*|*B*. When we attach 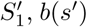 will become either *A*|(*B* ∪ *L*(*S*_2_)) or (*A* ∪ *L*(*S*_2_)) |*B*, depending on which side of *a*_1_ *s*′ is on.

Let *b*(*e*_1_) = *b*(*a*_1_) = *A*_*a*_|*B*_*a*_, so, when *e*′ is attached, edge *a*_1_ splits into two bipartitions: *A*_*a*_|(*B*_*a*_ ∪ *L*(*S*_2_)) and (*A*_*a*_ ∪ *L*(*S*_2_))|*B*_*a*_. If *s* = *e*_1_, then *b*(*s*) is one of these. If *s* ≠ *e*_1_, we know that *B*_*a*_ ⊂ *B*, because otherwise, *s* would lie between *e*_1_ and *e*, which is impossible by construction. Since *b*(*s*′) must be compatible with *A*_*a*_|(*B*_*a*_ ∪ *L*(*S*_2_)), this forces *b*(*s*′) to be *A*|(*B* ∪ *L*(*S*_2_)). Otherwise, neither side of (*A* ∪ *L*(*S*_2_)) | *B* is a subset of either side of *A*_*a*_ | (*B*_*a*_ ∪ *L*(*S*_2_)), making them incompatible. Thus, *b*(*s*) appears in *T* ′.

(2) Proof that GTM has *O*(*N* ^2^) worst case running time, where *N* is the number of taxa. The implementation of the GTM procedure can be summarized as shown in the algorithm block below.

#### Algorithm 1 GTM

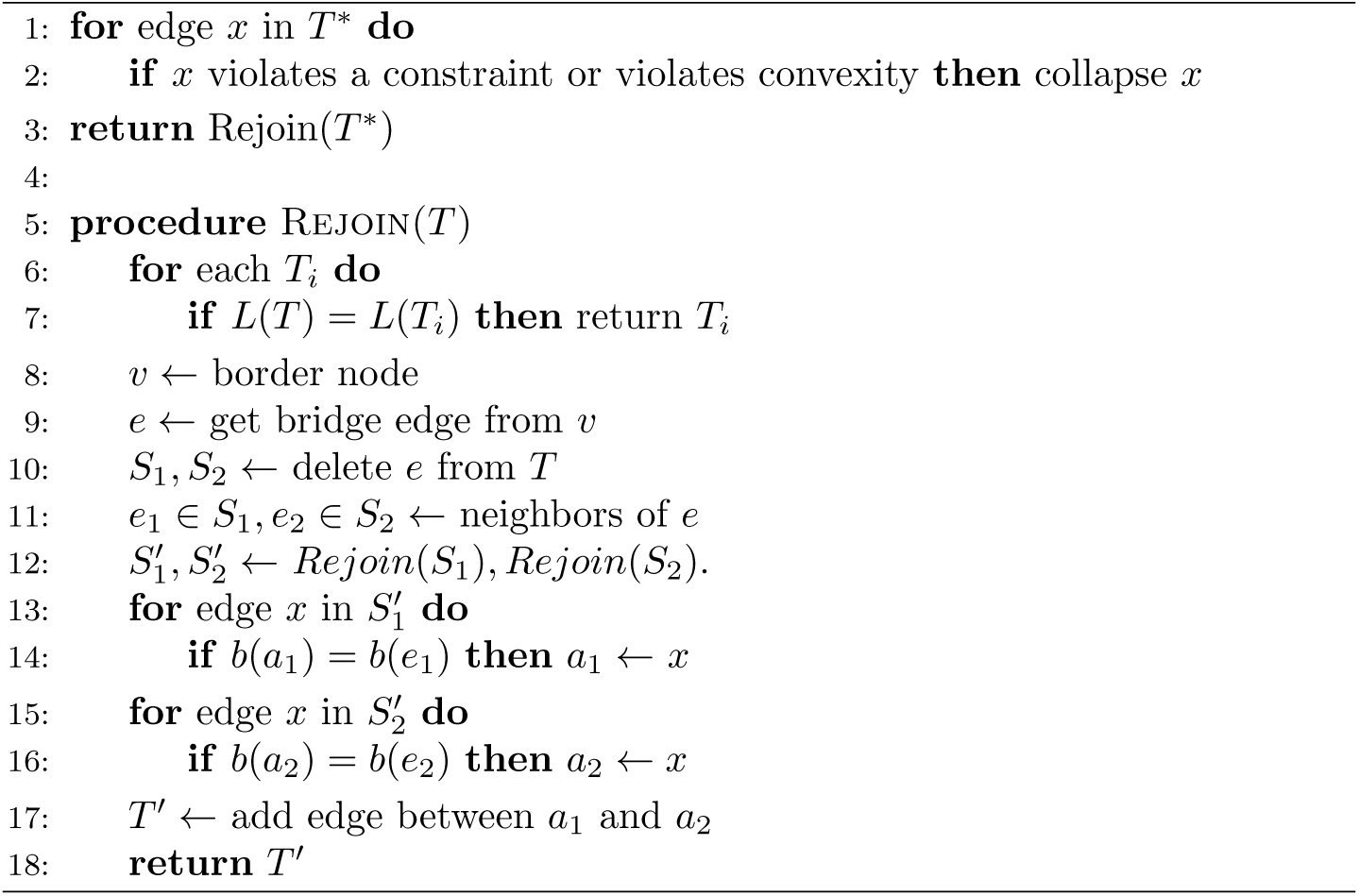

This permits a fairly straightforward runtime analysis. The first step is the collapse of violating edges in *T* *. In the interest of speed, we can facilitate this with a preprocessing step: for each constraint tree *T*_*i*_, we encode its bipartitions as binary bitmasks (in the conventional way) and store these numerical values in a hash table 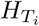. Then, for each edge *e* ∈ *T* * with associated bitmask *b*, we can check which constraint trees *e* satisfies by hashing the bits of *b* associated with each constraint tree and seeing if this hash appears in 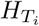. Since we’re hashing *N* bits for each of *N* edges in *T* *, this first part is 𝒪 (*N*^2^). While we’re traversing each edge, we also add some annotations to facilitate the second part of the algorithm quickly: based on which constraint tree each bit in *b* is associated with, we identify if *e* is a border edge separating which constraint trees, or which constraint set *e* lies in. This piggybacks on the total *O*(*N*) cost for *e*.

The second part is the recursive Rejoin itself. The recursion is invoked twice for each bridge edge, of which there are 𝒪(*k*), giving us 𝒪(*k*) calls to Rejoin. The check *L*(*T*) = *L*(*T*_*i*_) is *O*(*k*), since it uses the annotations we computed in the previous part. (We know which constraint trees were separated by the edge we just deleted to produce *T*). The next tasks are to find a border node *v*, as well as *a*_1_ and *a*_2_, and these are 𝒪(*N*) operations likewise facilitated by the computations from the previous part. Everything else in the procedure is a unit-time operation. Thus, we end up with 𝒪(*k*) calls of a 𝒪(*N*) function, so the second part is 𝒪(*Nk*). Putting everything together, the entire procedure has worst case running time 𝒪(*N*^2^).

### B.3 Proof of Theorem 3

**Theorem 3**: Let Φ be a model of evolution. If the method *X* used to construct the starting tree and the method *Y* used to construct the subset trees are both statistically consistent under Φ, then the DTM pipeline *X*-*Y* -GTM is also statistically consistent under Φ.

*Proof.* Let *T* be the model tree. Since *X* is statistically consistent, the starting tree produced by *X* will converge to the model tree, *T*, as the amount of data increases. When the starting tree is the model tree, the decomposition strategy (which is based on edge deletions from the starting tree) will produce subsets that are convex within the model tree. Since *Y* is also statistically consistent, the computed subset trees will also converge to the model species tree, restricted to each subset. Given correct subset trees, each of which is based on a convex subset within the model tree, the optimal unblended DTM-GT-FN tree is the model tree. Since GTM optimally solves unblended DTM-GT-FN, it will return *T* when given such an input. Hence, as the amount of data increases, *X*-*Y* -GTM will converge to the model tree *T*.

## C Supplementary Figures

**Figure 10:**
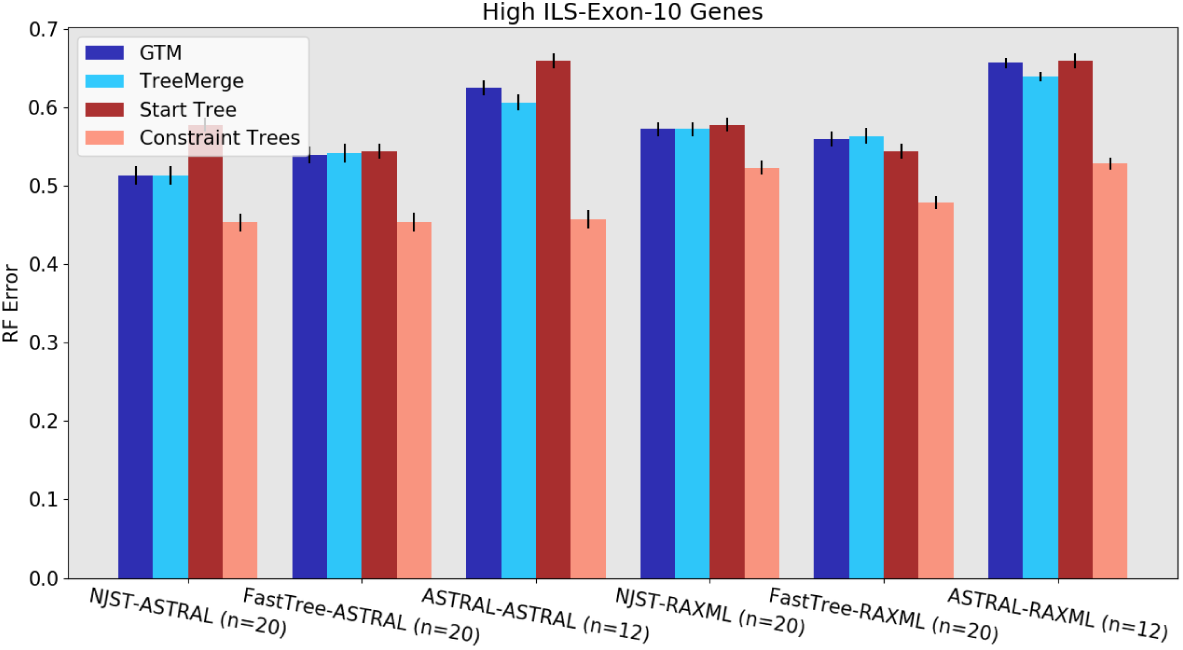
Experiment 1: Tree error rates for GTM and TreeMerge with different “starting tree-constraint tree” combinations over 10 high ILS exon genes with 1000 species. (NJMerge is not shown, because it failed for all high-ILS 10 and 25-gene replicates. Check Figure 12 for high-ILS NJMerge results). Both DTM methods work best with ASTRAL constraint trees and show a distinct preference for the NJst-ASTRAL combination. The value for *n* is the number of replicates being compared, where all methods ran. Error bars show standard error of the replicate average.

**Figure 11:**
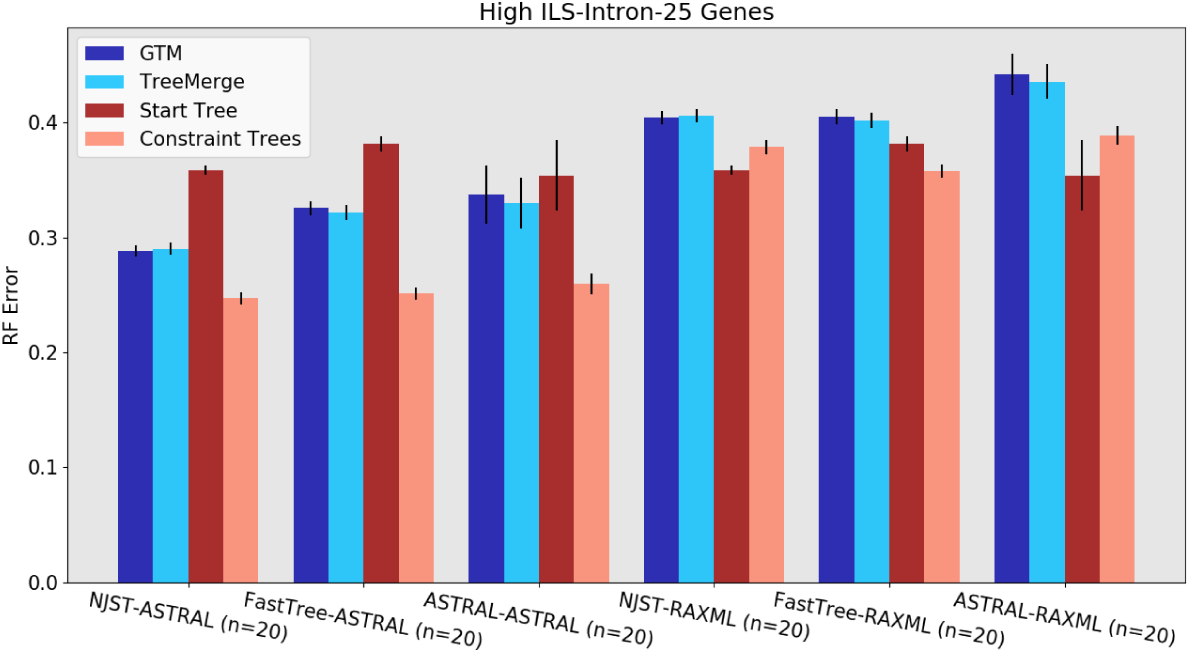
Experiment 1: Tree error rates for GTM and TreeMerge with different “starting tree-constraint tree” combinations over 25 high ILS intron genes with 1000 species. (NJMerge is not shown, because it failed for all high-ILS 10 and 25-gene replicates. Check Figure 12 for high-ILS NJMerge results). Both DTM methods work best with ASTRAL constraint trees and show a distinct preference for the NJst-ASTRAL combination. The value for *n* is the number of replicates being compared, where all methods ran. Error bars show standard error of the replicate average.

**Figure 12:**
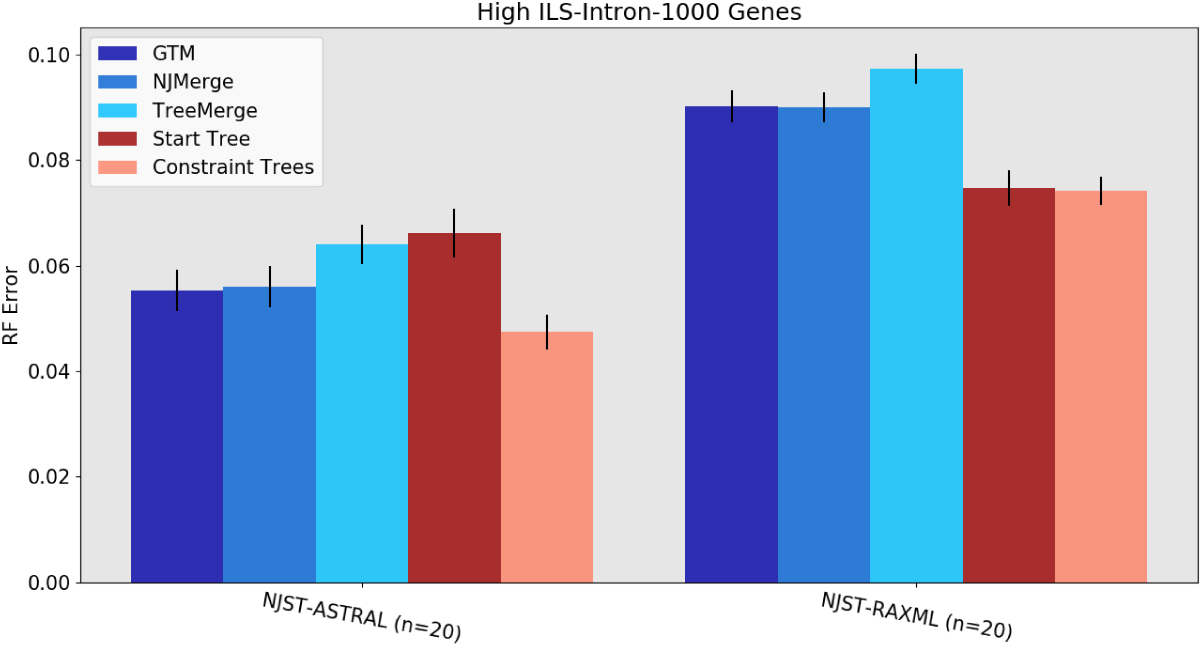
Experiment 1: Tree error rates for GTM, NJMerge, and TreeMerge with NJst starting trees and ASTRAL/RAxML constraint trees over 1000 high ILS intron genes with 1000 species. Only NJst starting trees are used, due to impracticability of FastTree and ASTRAL starting trees at this scale. All three DTM methods work best with ASTRAL constraint trees. The value for *n* is the number of replicates being compared, where all methods ran. Error bars show standard error of the replicate average.

**Figure 13:**
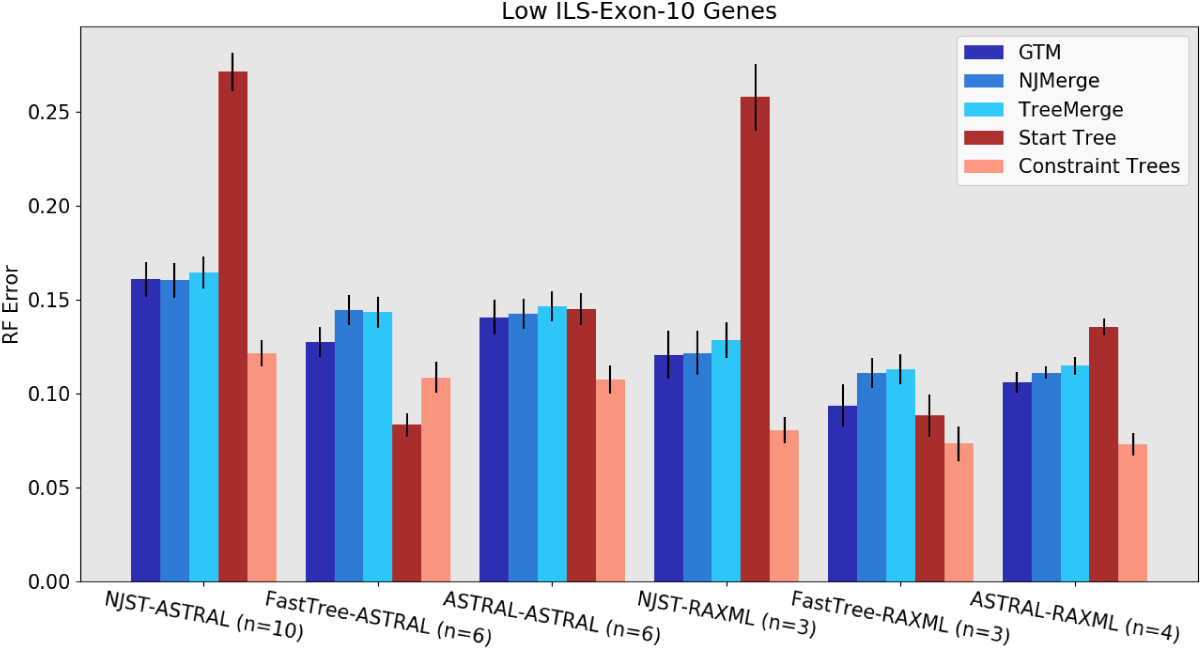
Experiment 1: Tree error rates for GTM, NJMerge, and TreeMerge with different “starting tree-constraint tree” combinations over 10 low ILS exon genes with 1000 species. All three DTM methods work best with FastTree starting trees and RAxML constraint trees. The value for *n* reflects the the number of replicates being compared, where all methods ran. Error bars show standard error of the replicate average.

**Figure 14:**
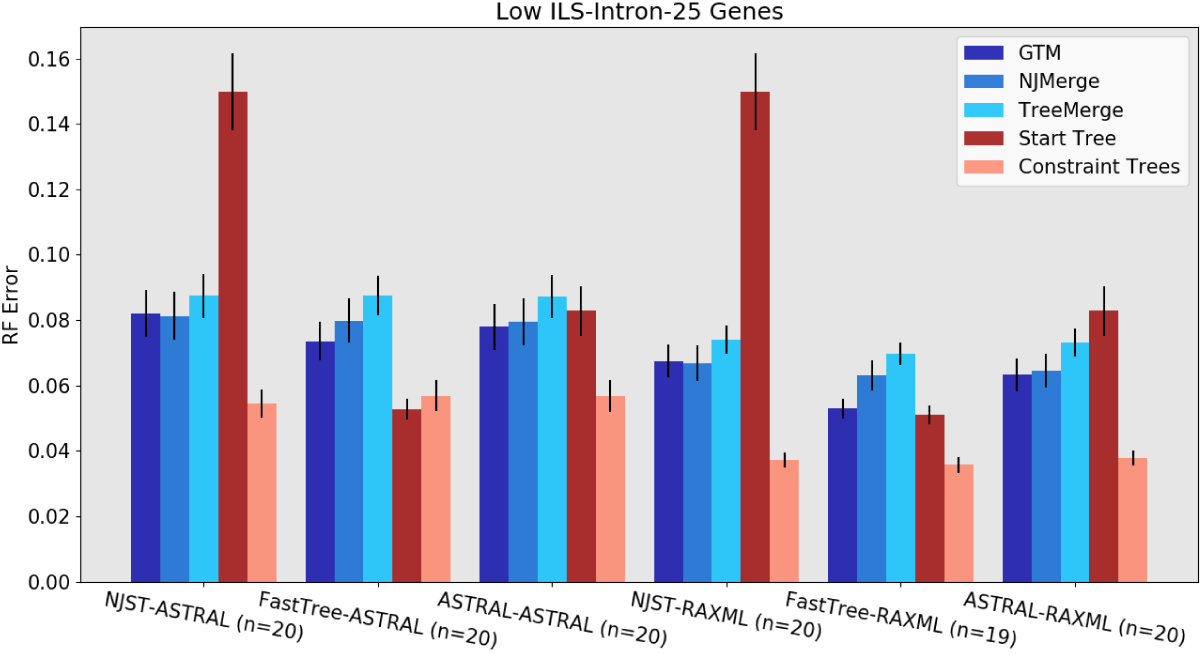
Experiment 1: Tree error rates for GTM, NJMerge, and TreeMerge with different “starting tree-constraint tree” combinations over 25 low ILS intron genes with 1000 species. All three DTM methods work best with FastTree starting trees and RAxML constraint trees. The value for *n* is the number of replicates being compared, where all methods ran. Error bars show standard error of the replicate average.

**Figure 15:**
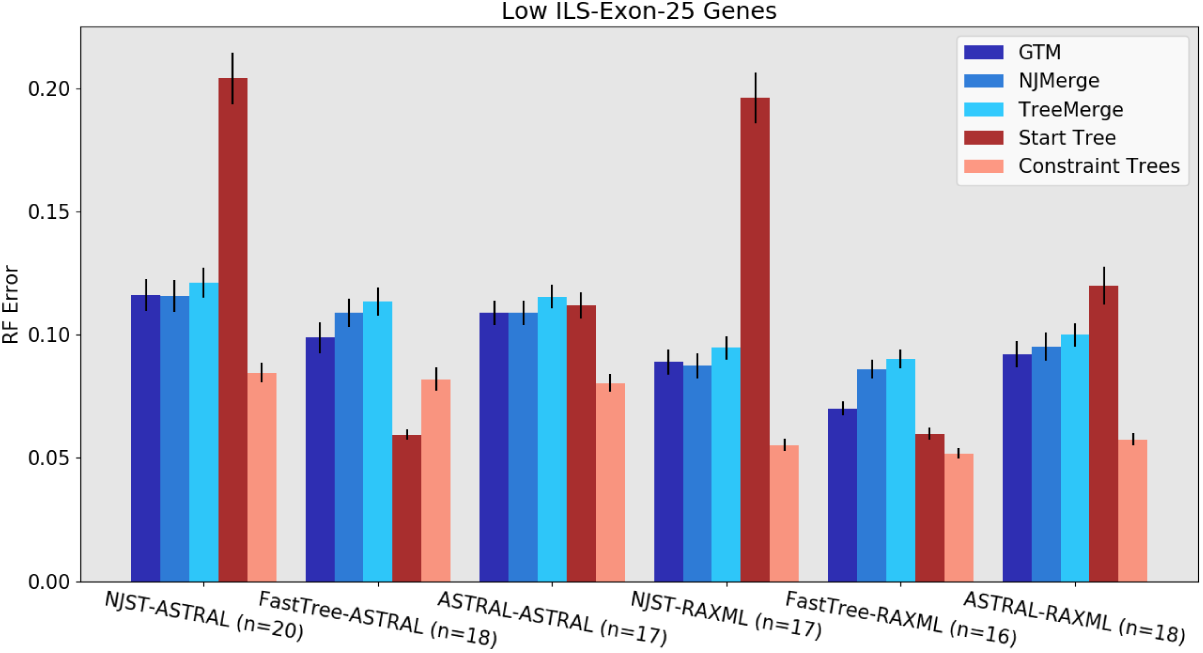
Experiment 1: Tree error rates for GTM, NJMerge, and TreeMerge with different “starting tree-constraint tree” combinations over 25 low ILS exon genes with 1000 species. All three DTM methods work best with RAxML constraint trees. All three methods improve with the FastTree starting tree, with a big improvement for GTM. The value for *n* is the number of replicates being compared, where all methods ran. Error bars show standard error of the replicate average.

**Figure 16:**
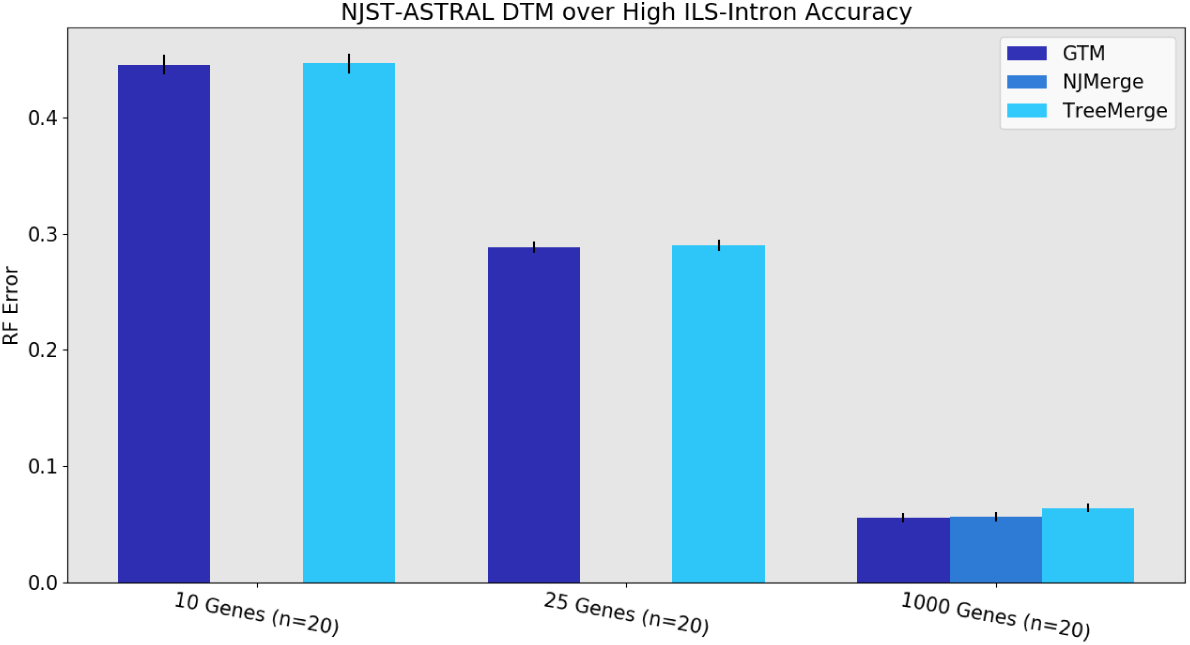
Experiment 2: Tree error rates for three DTM methods on 1000 species given NJst starting tree and ASTRAL constraint trees, the best pipeline for high ILS from Experiment 1. NJMerge failed to complete within the allowed 4 hour period for all 10- and 25-gene high ILS replicates (results shown for NJMerge on 1000-gene datasets are from [3]). GTM and TreeMerge are about even, with a slight advantage to GTM at 1000 genes. The value for *n* is the number of replicates being compared where all methods ran. Error bars show standard error of the replicate average.

**Figure 17:**
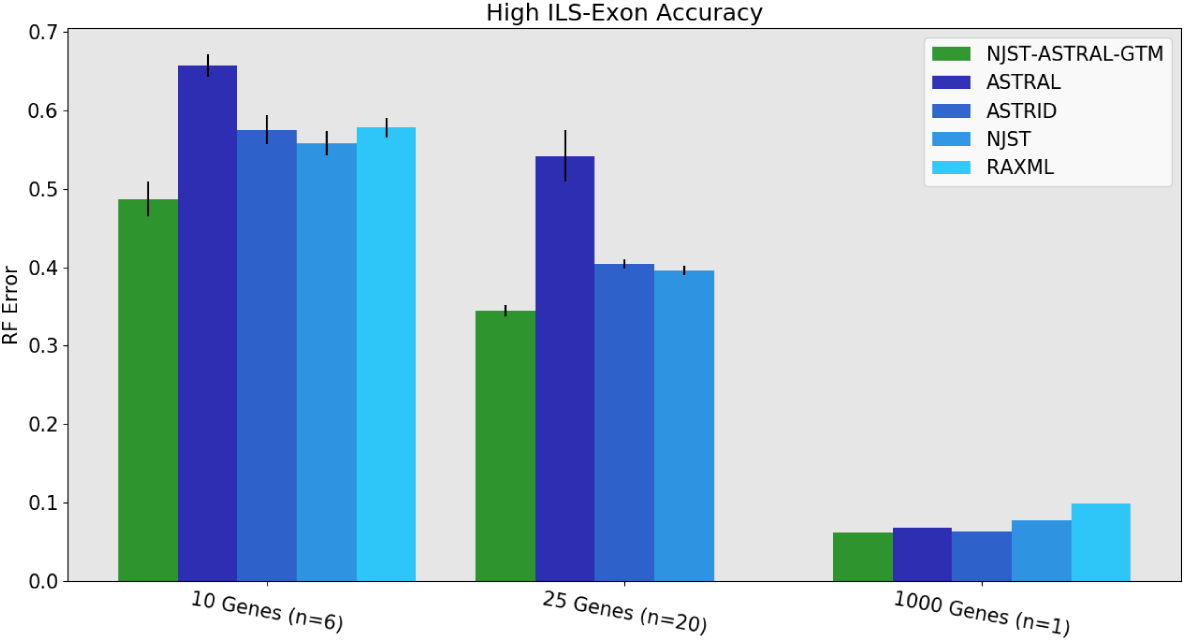
Experiment 3: Tree error rates for NJst-ASTRAL-GTM, RAxML, and leading summary methods (ASTRAL, ASTRID, NJst) on high ILS datasets with 1000 species. GTM is more accurate than the other methods on small numbers of genes, and roughly matches ASTRAL and ASTRID on 1000. The value for *n* is the number of replicates being compared, where all methods ran (i.e., where ASTRAL and RAxML are both available). RAxML timed out on 8 of the 10-gene replicates and all 25-gene replicates. ASTRAL timed out on 7 10-gene replicates. The 1000-gene ASTRAL and RAxML trees were taken from the NJMerge study [3], and only one ASTRAL tree is available for high ILS exons. Error bars indicate standard error of the replicate average.

**Figure 18:**
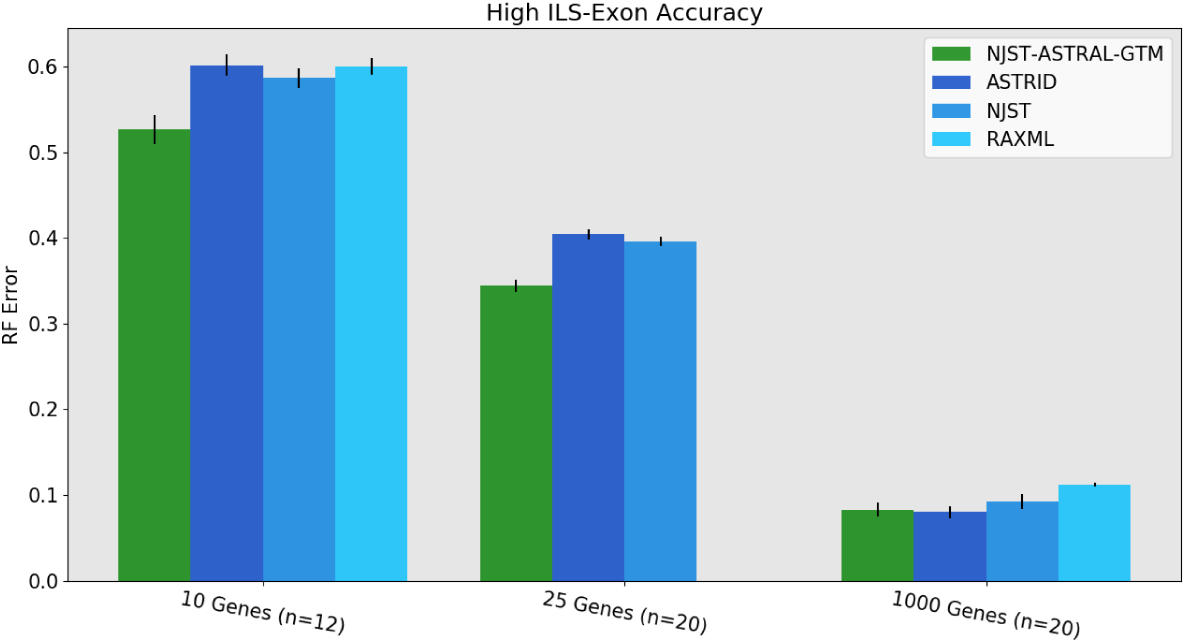
Experiment 3: Tree error rates for NJst-ASTRAL-GTM, RAxML, and summary methods (ASTRID and NJst) on high ILS datasets with 1000 species. NJst-ASTRAL-GTM is more accurate than the other methods on small numbers of genes. For 1000-gene datasets, NJst-ASTRAL-GTM and ASTRID are tied, and are more accurate than NJst and RAxML. The value for *n* is the number of replicates where RAxML trees are available. RAxML timed out on 8 of the 10-gene replicates and all 25-gene replicates. The 1000-gene RAxML trees were taken from the NJMerge study [3]. Error bars show standard error of the replicate average.

**Figure 19:**
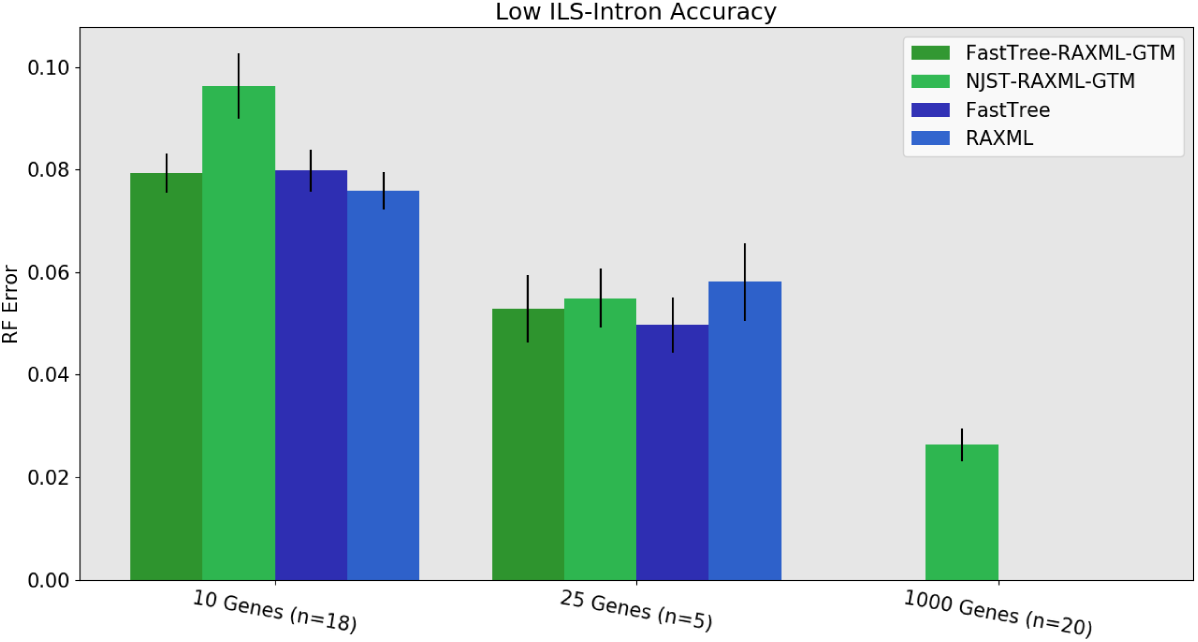
Experiment 3: Comparison of FastTree-RAxML-GTM, NJst-RAxML-GTM, FastTree, and RAxML on 1000-species datasets with low ILS introns. The value for *n* is the number of replicates on which RAxML completed; missing replicates indicate RAxML exceeding runtime limits on 10 and 25 genes, and RAxML trees were not available for 1000 genes. FastTree was not used for 1000 genes. Error bars show standard error of the replicate average.

**Figure 20:**
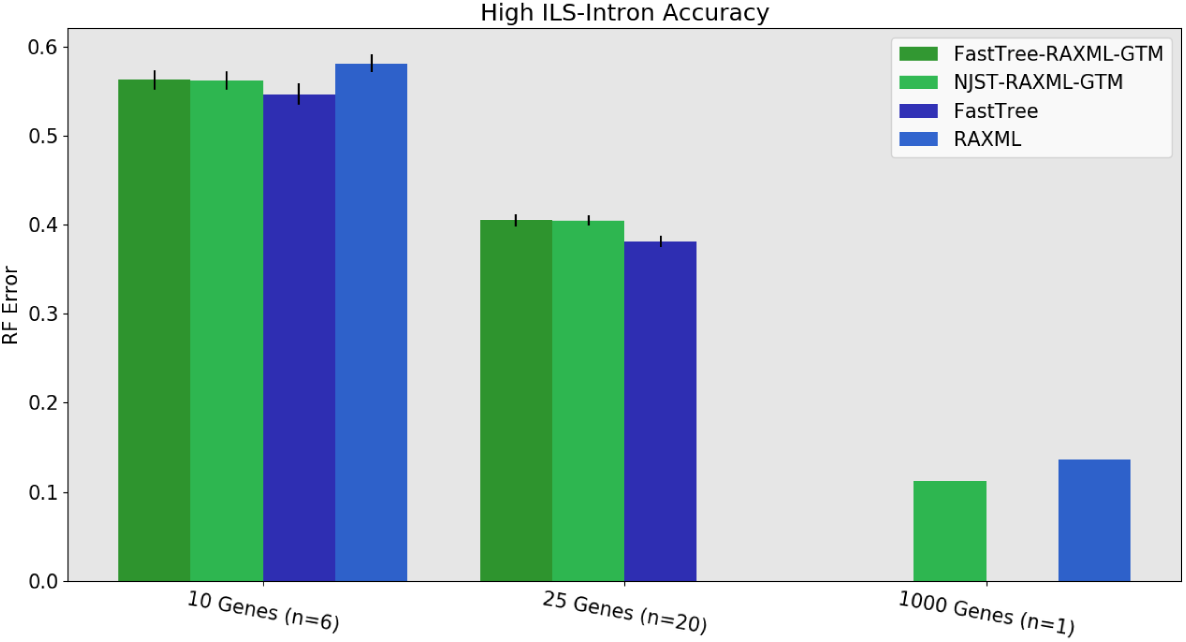
Experiment 3: Comparison of FastTree-RAxML-GTM, NJst-RAxML-GTM, FastTree, and RAxML on 1000-species datasets with high ILS introns. The value for *n* is the number of replicates on which RAxML completed; missing replicates indicate RAxML exceeding runtime limits on 10 and 25 genes (the 1000-gene RAxML trees are taken from [3]). FastTree was not used for 1000 genes. Error bars show standard error of the replicate average.

**Figure 21:**
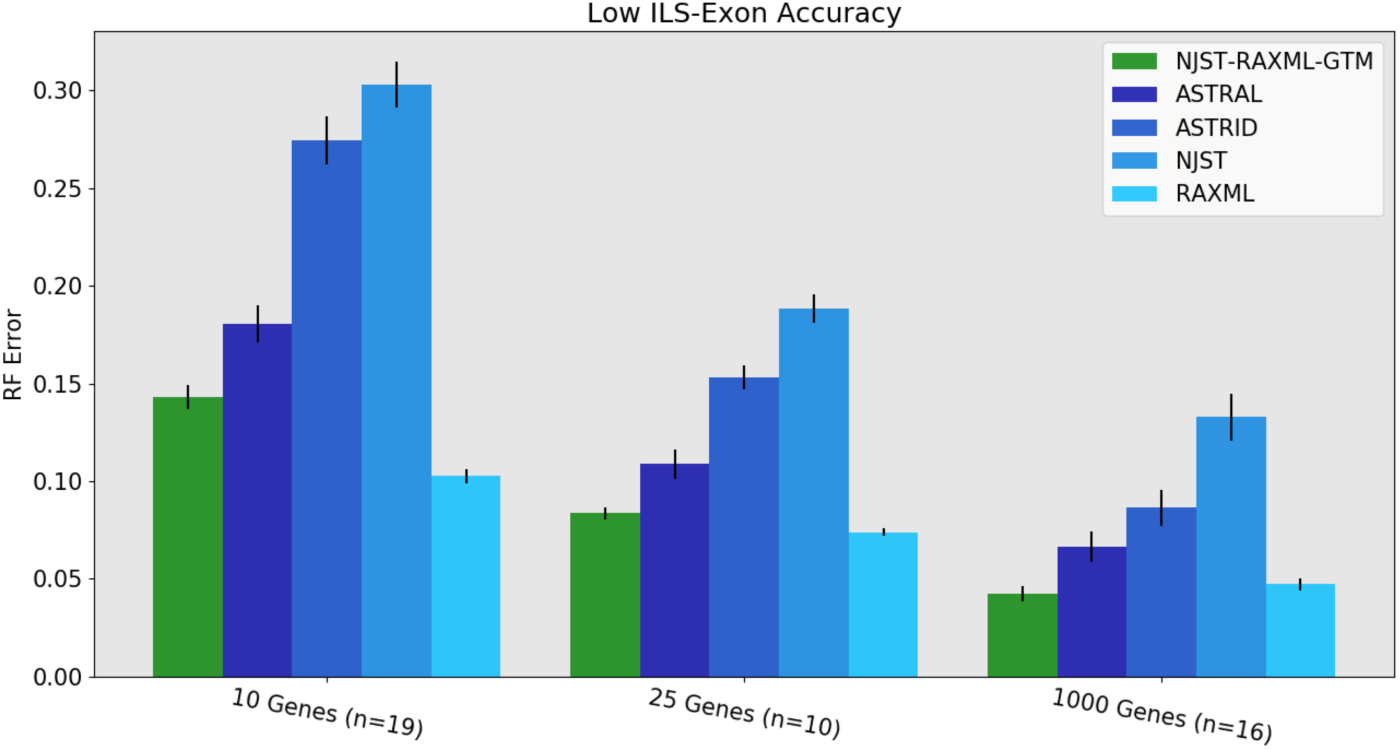
Experiment 3: Comparison of NJst-RAxML-GTM to RAxML and other species tree methods on 1000-species datasets with low ILS. The value for *n* is the number of replicates on which all 5 methods completed; missing replicates indicate RAxML exceeding runtime limits on 10 and 25 genes (the 16 1000-gene RAxML trees are taken from [3]). Error bars show standard error of the replicate average

## D Supplementary Tables

**Table 7:** Number of replicates, out of 20 in each model condition, that RAxML, ASTRAL, and NJMerge complete on within the allowed 4 hour timeframe on 1000-species datasets.

|  | RAxML | ASTRAL | NJMerge |
| --- | --- | --- | --- |
| 1000 exons, high ILS | 0 | 1 | 20 |
| 1000 introns, high ILS | 0 | 16 | 20 |
| 1000 exons, low ILS | 0 | 20 | 20 |
| 1000 introns, low ILS | 0 | 20 | 20 |
| 25 exons, high ILS | 0 | 20 | 0 |
| 25 introns, high ILS | 0 | 20 | 0 |
| 25 exons, low ILS | 10 | 20 | 17 |
| 25 introns, low ILS | 5 | 20 | 20 |
| 10 exons, high ILS | 12 | 13 | 0 |
| 10 introns, high ILS | 6 | 19 | 0 |
| 10 exons, low ILS | 19 | 20 | 3 |
| 10 introns, low ILS | 18 | 20 | 14 |

**Table 8:** Tree error rates for NJst-RAxML-GTM and RAxML on 1000-species datasets with low ILS. The value for *n* is the number of replicates being compared, where a RAxML tree is available (finished within the time limit). The 16 exon 1000-gene RAxML trees are taken from the NJMerge study. Improved accuracy can be obtained using FastTree-RAxML-GTM.

|  | NJst-RAxML-GTM | RAxML |
| --- | --- | --- |
| 10 Exons (n=19) | 0.143 | 0.103 |
| 10 Introns (n=18) | 0.096 | 0.076 |
| 25 Exons (n=10) | 0.083 | 0.074 |
| 25 Introns (n=5) | 0.055 | 0.058 |
| 1000 Exons (n=16) | 0.042 | 0.047 |
| 1000 Introns (n=20) | 0.026 | N/A |

**Table 9:** Tree error rates for high ILS, NJst-ASTRAL-GTM vs ASTRAL. The value for *n* is the number of replicates being compared (where ASTRAL finished within the time limit). 1000 species, max subset size 120.

|  | NJst-ASTRAL-GTM | ASTRAL |
| --- | --- | --- |
| 10 Exons (n=13) | 0.489 | 0.663 |
| 10 Introns (n=19) | 0.442 | 0.646 |
| 25 Exons (n=20) | 0.344 | 0.542 |
| 25 Introns (n=20) | 0.289 | 0.354 |
| 1000 Exons (n=1) | 0.062 | 0.067 |
| 1000 Introns (n=16) | 0.057 | 0.059 |

**Table 10:** Average runtime (seconds) over low ILS exons, NJst-RAxML-GTM vs RAxML on 1000-species datasets. The value for *n* is the number of replicates being compared, i.e., where a RAxML tree is available. The 1000-gene RAxML trees are taken from the NJMerge study, and these are RAxML’s last result on a 48-hour time limit (hence, the flat 172800 second timing for these). The RAxML constraint trees for the 1000-gene NJst-RAxML-GTM analyses were also used from the NJMerge study [3], and timings are shown for a subset of replicates that were rerun to gather timings. Finally, the 1000 FastTree2 gene trees used for NJst were reported to take 65 minutes. This is scaled to the number of genes being used.

|  | NJst-RAxML-GTM | RAxML |
| --- | --- | --- |
| **10 Genes (n=19)** |  |  |
| -Gene trees | 39.0 | n.a. |
| -NJst | 7.6 | n.a. |
| -RAxML subtrees | 864.3 | n.a. |
| -GTM | 0.4 | n.a. |
| -Total | 911.3 | 7,313.7 |
| **25 Genes (n=10)** |  |  |
| -Gene trees | 98.0 | n.a. |
| -NJst | 7.6 | n.a. |
| -RAxML subtrees | 1,504.8 | n.a. |
| -GTM | 0.4 | n.a. |
| -Total | 1,610.8 | 10,539.4 |
| **1000 Genes (n=17)** |  |  |
| -Gene trees | 3,900.0 | n.a. |
| -NJst | 7.7 | n.a. |
| -RAxML subtrees | 69,322.4 | n.a. |
| -GTM | 0.4 | n.a. |
| -Total | 73,230.5 | 172,800.0 |

**Table 11:**
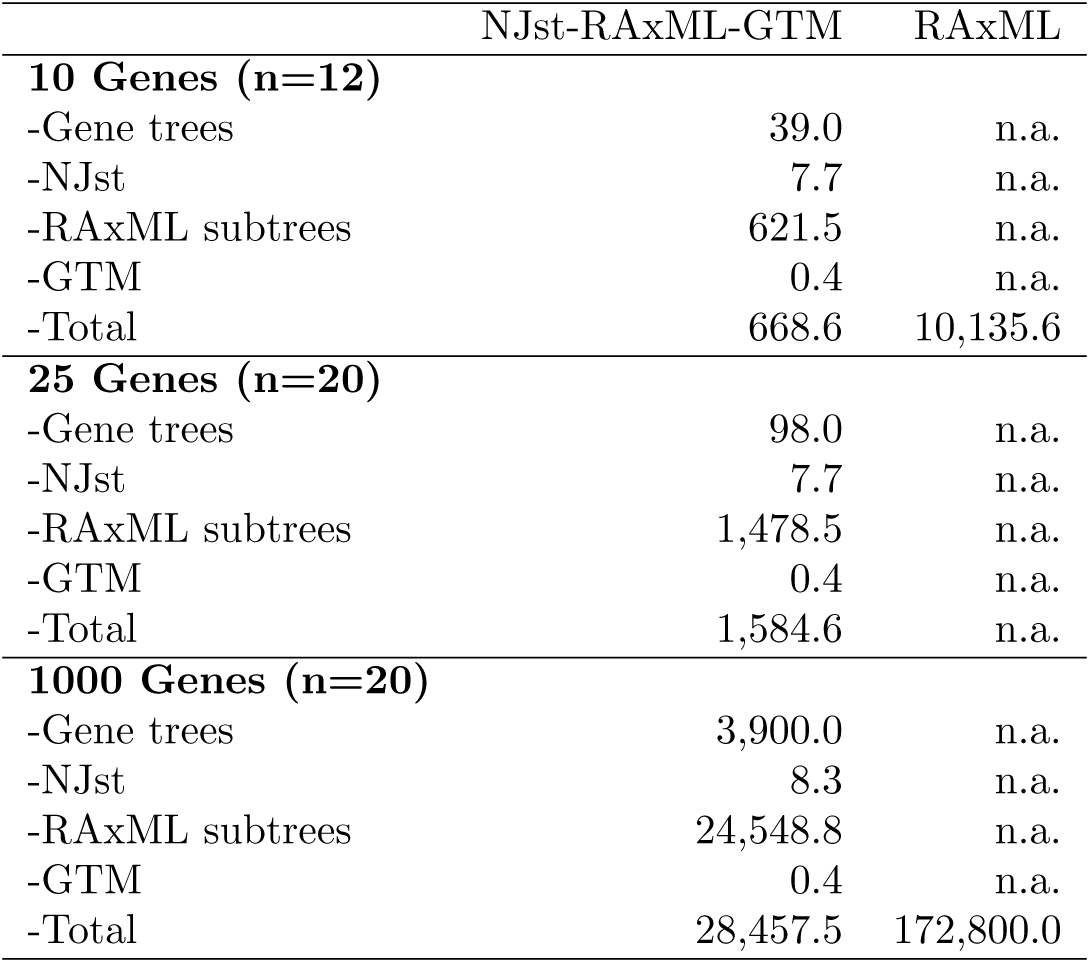
Average runtime (seconds) of NJst-RAxML-GTM and RAxML on 1000-species high ILS datasets with varying numbers of exons. The value for *n* is the number of replicates being compared, i.e., where a RAxML tree is available. The 1000-gene RAxML trees are taken from the NJMerge study, and these are RAxML’s last result on a 48-hour time limit (hence, the flat 172800 second timing for these). The RAxML constraint trees for 1000-gene NJst-RAxML-GTM were also used from the NJMerge study [3], and timings are shown for a subset of replicates that were rerun to gather timings. Finally, the 1000 FastTree2 gene trees used for NJst were reported to take 65 minutes. This is scaled to the number of genes being used.

|  | NJst-RAxML-GTM | RAxML |
| --- | --- | --- |
| **10 Genes (n=12)** |  |  |
| -Gene trees | 39.0 | n.a. |
| -NJst | 7.7 | n.a. |
| -RAxML subtrees | 621.5 | n.a. |
| -GTM | 0.4 | n.a. |
| -Total | 668.6 | 10,135.6 |
| **25 Genes (n=20)** |  |  |
| -Gene trees | 98.0 | n.a. |
| -NJst | 7.7 | n.a. |
| -RAxML subtrees | 1,478.5 | n.a. |
| -GTM | 0.4 | n.a. |
| -Total | 1,584.6 | n.a. |
| **1000 Genes (n=20)** |  |  |
| -Gene trees | 3,900.0 | n.a. |
| -NJst | 8.3 | n.a. |
| -RAxML subtrees | 24,548.8 | n.a. |
| -GTM | 0.4 | n.a. |
| -Total | 28,457.5 | 172,800.0 |

**Table 12:** Average runtime (seconds) over high ILS introns on 1000-species datasets for NJst-ASTRAL-GTM vs. ASTRAL. The value for *n* is the number of replicates being compared, where ASTRAL trees are available. The 1000-gene ASTRAL trees are taken from the NJMerge study [3]. The 1000 FastTree2 gene trees were reported to take 65 minutes. This is scaled to the number of genes being used and applied to both methods. 1000 species, max subset size 120.

|  | NJst-ASTRAL-GTM | ASTRAL |
| --- | --- | --- |
| **10 Genes (n=18)** |  |  |
| -Gene trees | 39.0 | 39.0 |
| -NJst | 7.7 | n.a. |
| -ASTRAL | 50.7 | 8,617.0 |
| -GTM | 0.4 | n.a. |
| -Total | 97.8 | 8,656.0 |
| **25 Genes (n=20)** |  |  |
| -Gene trees | 98.0 | 98.0 |
| -NJst | 7.7 | n.a. |
| -ASTRAL | 69.0 | 5,441.4 |
| -GTM | 0.4 | n.a. |
| -Total | 175.1 | 5,539.4 |
| **1000 Genes (n=16)** |  |  |
| -Gene trees | 3,900.0 | 3,900.0 |
| -NJst | 7.7 | n.a. |
| -ASTRAL | 4,048.9 | 149,145.9 |
| -GTM | 0.4 | n.a. |
| -Total | 7,949.3 | 153,045.9 |

